# Landscape structure and dispersal rate drive large scale catastrophic shifts in spatially explicit metapopulations

**DOI:** 10.1101/2021.11.19.469221

**Authors:** Camille Saade, Emanuel A. Fronhofer, Benoît Pichon, Sonia Kéfi

## Abstract

Even when environments deteriorate gradually, ecosystems may shift abruptly from one state to another. Such catastrophic shifts are difficult to predict and reverse (hysteresis). While well studied in simplified contexts, we lack a general understanding of how catastrophic shifts spread in realistic spatial contexts. For different types of landscape structure, including typical terrestrial modular and riverine dendritic networks, we here investigate landscape-scale stability in metapopulations made of bistable patches. We find that such metapopulations usually exhibit large scale catastrophic shifts and hysteresis, and that the properties of these shifts depend strongly on metapopulation spatial structure and dispersal rate: intermediate dispersal rates and a riverine spatial structure can largely reduce hysteresis size. Interestingly, our study suggests that large-scale restoration is easier with spatially clustered restoration efforts and in populations characterized by an intermediate dispersal rate.

## Introduction

As humans exert increasing pressures on ecosystems, understanding how they respond in terms of biomass, species diversity and composition is one of the most pressing issue in ecology. Importantly, some ecosystems can exhibit abrupt responses to gradual changes in environmental conditions (Noy-Meir, 1975; May, 1977; Scheffer et al., 2001). Such responses, referred to as “catastrophic shifts”, are usually attributed to the existence of multiple stable ecosystem states between which ecosystems can shift following a perturbation or when a threshold in environmental condition, or “tipping point” is passed (Suding et al., 2004; Kéfi et al., 2007b, 2016). Reverting the ecosystem to its initial state after a shift can be challenging if not impossible, a phenomenon known as “hysteresis”. Catastrophic shifts have been described in various ecosystems all over the world (Biggs et al., 2018). Common examples include the eutrophication of shallow lakes (Scheffer et al., 1993; Meijer et al., 1994; Scheffer et al., 1997; Carpenter et al., 1999; Jeppesen et al., 1999), the degradation of coral reefs (Done, 1992; Knowlton, 1992; McCook, 1999; Nyström et al., 2000), the transitions between woodlands and savannas (Dublin et al., 1990; Wilson and Agnew, 1992; Walker, 1995) and the desertification of grasslands (Kassas, 1995; Rietkerk et al., 1997; Nicholson, 2000; Wang and Eltahir, 2000; Reynolds et al., 2007; Kéfi et al., 2007a). Because they are difficult to predict and reverse (Scheffer and Carpenter, 2003) and because they can negatively affect human livelihoods (Reynolds et al., 2007; Biggs et al., 2018), catastrophic shifts have gained a large interest in the literature, providing us with a theoretical framework rooted in bifurcation theory to describe the most common catastrophic shifts (*e.g*., desertification: Noy-Meir 1975; May 1977; Rietkerk and van de Koppel 1997; Klausmeier 1999; Kéfi et al. 2007b,a; or shallow lakes eutrophication: Scheffer et al. 2001; Carpenter et al. 1999).

However, early theoretical work on catastrophic shifts mainly focused on isolated systems (but see Keitt et al., 2001; van Nes and Scheffer, 2005). Real ecosystems are networks of connected entities between which matter and energy can flow (Leibold et al., 2004). This means that if one of these entities experiences a catastrophic shift locally, this shift can spread through the spatial network and possibly trigger other shifts. Scaling up our understanding of catastrophic shifts from the local scale to the regional scale remains a challenge. In particular, whether local multistability implies regional multistability or whether different stable states can coexist in space (spatial multistability) and how this is modulated by a landscape’s spatial structure are still open questions. Recent studies have started to address these questions by including space explicitly in models, under two main settings: *i*) in continuous space, which is well suited to relatively homogeneous habitats where clear spatial entities are not easily identified (see the example of Lake Veluwe in van de Leemput et al., 2015), and *ii*) in discrete space, better representing habitats with clear spatial entities (or patches) connected through dispersal of individuals and flows of resources (e.g., an archipelago such as the Åland Islands in Hanski et al., 1995). As a consequence of these recent developments, we have a good understanding of how locally bistable dynamics can affect the regional behaviour of spatially structured ecosystems: alternative stable states are not expected to coexist in space (Keitt et al., 2001; van de Leemput et al., 2015), unless dispersal is low and space is discrete (Keitt et al., 2001), or strong stochasticity and heterogeneity are at play (van Nes and Scheffer, 2005; Martín et al., 2015). A spatially structured system will experience sharp transitions between fully occupied and fully empty states. These can usually not be called catastrophic shifts as regional bistability and hysteresis are largely reduced due to the dominance of a single of the two possible stable states (Keitt et al., 2001; Hilt et al., 2011; van de Leemput et al., 2015). However, all studies so far assume overly simplified spatial structures, either 1-D (Keitt et al., 2001; van de Leemput et al., 2015) or 2-D continuous space (Martín et al., 2015), linear (Keitt et al., 2001; Hilt et al., 2011) or grid-like (van Nes and Scheffer, 2005) discrete systems. This implies that we currently don’t know how more realistic spatial settings affect landscape-scale stability. Indeed, in real ecosystems, suitable habitat is usually neither continuous nor organized on a regular lattice but has a particular structure: terrestrial populations, for example, usually show emergent modularity (Gilarranz, 2020) while riverine systems are typically dendritic (Carraro et al., 2020; Rinaldo et al., 2020). Importantly, these specific spatial configurations have been shown to affect ecological dynamics (Gilarranz et al., 2017; Altermatt and Fronhofer, 2018; Carrara et al., 2012). This omission of spatial complexity is an important shortcoming of the current state of the literature as the properties of such habitats may change how local bistability affects regional scale dynamics and equilibria (van Nes and Scheffer, 2005; van de Leemput et al., 2015).

Here, we aim to fill this gap by studying how local bistability affects the regional scale dynamics of spatially complex populations, using a metapopulation framework in which local patches are connected to each other via dispersal of individuals. We simulate the dynamics of metapopulations with various spatial structures, ranging from classical abstract structures (lattice and random, Erdős–Rényi, spatial networks) to structures rooted in real systems (Random Geometric Graphs for terrestrial systems and Optimal Channel Networks for riverine systems; Gilarranz, 2020; Carraro et al., 2020). For a range of dispersal rates, we measure the size of hysteresis and the position of tipping points at the scale of the whole landscape. We also study how the structure of perturbations themselves affects landscape dynamics by conducting targeted local perturbations of firstly high-degree patches *vs*. low degree patches and secondly neighboring patches *vs*. independent patches.

## Model description

### Metapopulation model

We used a model of a spatially structured metapopulation describing the dynamics of *n* patches linked by the dispersal of individuals. Here is a short description of the model, but see the supplement S1 for a more in-depth presentation. The dynamics of the vector 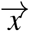 (containing the local biomasses *x_i_*) are described by a system of *n* coupled differential equations, which can be written in a matricial form:

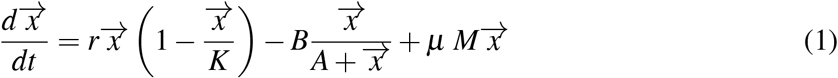

The first two terms of Eq. 1 describe the local dynamics of the biomass, while the last term describes the dispersal between patches.

#### Local dynamics

We describe the local dynamics using a model derived by Noy-Meir (1975) in which plants grow logistically with a growth rate *r* and carrying capacity *K*, and are harvested by a fixed consumer with a Holling type II functional response (*B*: maximal harvesting rate, *A*: half saturation constant). For the rest of this work, we used the same fixed values for *r, K* and *A* (Table S6.1). These parameter values give rise to two tipping points at *B* = 50 and *B* = 56.25 between which the system is bistable: it admits two stable equilibria (*x_i_* = 0 and *x_i_* = *r*_1_ > 0) separated by an unstable equilibrium (0 < *r*_2_ < *r*_1_) (Fig. 1a).

**Figure 1:**
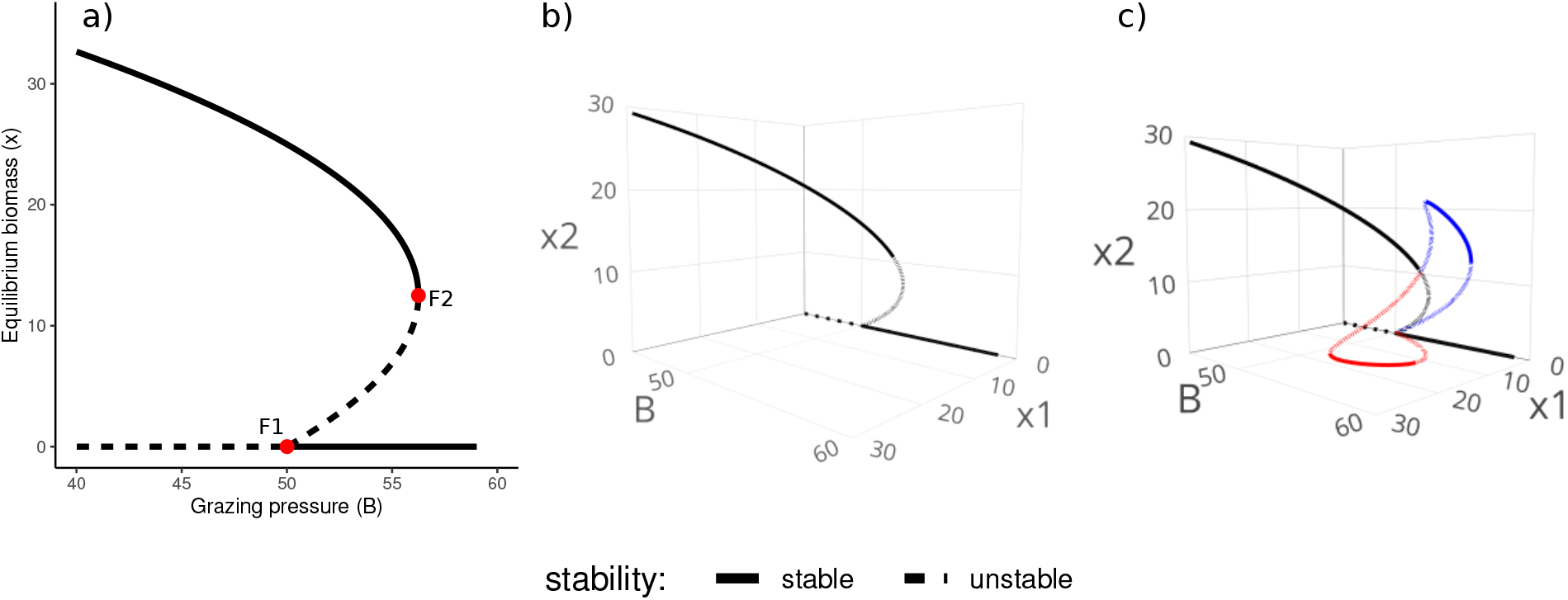
Bifurcation diagrams of a single patch (a) and of two patch systems (b, c) at high (b, *μ* = 0.3) and low (c, *μ* = 0.005) dispersal rates. Full lines are stable equilibria, dashed lines are unstable equilibria. (a) Equilibria of the biomass (*x*) of an isolated patch as a function of the grazing pressure (*B*). (b, c) Equilibria of the biomasses (*x*_1_, *x*_2_) of two patches connected by dispersal, as a function of the grazing pressure (*B*).

Note that this is one of several possible models of bistability that we chose because of its simplicity and intuitive biological interpretation. Importantly, we show in the supplement S3 how this model and three other common models of bistable systems share a common structure through parameter aggregation which highlights the generality of our findings.

#### Spatial dynamics

We consider that individuals leave their local patch at a constant rate *μ* and that the dispersing individuals are distributed evenly among adjacent patches. Mathematically, the vector describing net dispersal for all patches is given by the last term of Eq. 1 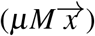 where *M* is a matrix describing the structure of the landscape: the diagonal terms describe emigration (*M_i,i_* = −1) and the off-diagonal terms describe the neighbourhood relationship between patches (see detail in supplement S1).

#### Integration scheme

We simulated the dynamics using the function *ode* (R-package deSolve, version 1.28). We performed linear regressions on the biomass of each patch over the last 100 simulated values to determine whether the metapopulation had reached an equilibrium (slope smaller than 10^−3^).

### Analysis of a two-patch system

Before moving on to large systems which are only accessible through simulation (see section “Landscape design”), we conducted analytically the stability analysis of a simple system for demonstrating general principles. We used a particular case of Eq. 1 with only two patches exchanging biomass at a rate *μ* (supplement S1: “Two-patch system”). We determined all its existing equilibria as well as their stability using *MatCont* (version 7.2, in *MATLAB R2017b*).

### Landscape design

We used four different types of landscapes (see supplement S2 for more details on landscape generation, and Fig. S6.1 to S6.3 for examples of landscapes): *i*) Regular graphs, where patches are arranged regularly on a lattice with periodic boundary conditions. While not very realistic, such landscapes have been used in previous studies (Keitt et al., 2001; van Nes and Scheffer, 2005) and we include them for the sake of comparison. *ii*) Erdös-Rényi graphs are obtained by randomly connecting pairs of patches with a fixed probability *p* and can be thought of as a null model. *iii*) Random Geometric Graphs (RGGs) are obtained by randomly drawing patch coordinates in space and connecting patches depending on distance. The random distribution of patches in space results in the emergence of modularity which is thought to be characteristic of terrestrial systems (Gilarranz, 2020). *iv*) Optimal Channel Networks (OCNs) are generated by simulating geomorphological processes to obtain a structure closely resembling that of a river (Carraro et al., 2020).

For each type of landscape, we generated 50 replicates with each *n* = 100 patches. The main text analysis was made on landscapes where patches had on average approximately 4 neighbours (except for OCNs as their generating process constrains their connectivity), and we conducted sensitivity analysis on the connectivity using networks with higher and lower connectivity (supplement S4).

### Simulations

For each type of network, we generated 50 replicates and computed their bifurcation diagram for 100 values of dispersal rates (*μ*) between 0.001 and 1. We used 46 values of harvesting rates (*B*) equally spaced between 50 and 56.25 (the bistability range of a single patch). For each value of *B*, we computed the high-biomass (resp. low-biomass) branch of the bifurcation diagram by initially setting each patch to the positive (resp. null) equilibrium of a single patch. Once the system had reached an equilibrium, we degraded (resp. restored) 5% of the patches by setting them to a null biomass (resp. to their positive equilibrium). We then measured the steady-state average biomass in order to plot the bifurcation diagram.

We used three different perturbation schemes: Firstly, we chose the 5 patches to be degraded/restored randomly (Fig. 2–4). Secondly, we focused on the influence of node degree by degrading/restoring patches chosen from either the most or least connected nodes (Fig. 5). We conducted these perturbations in Erdös-Rényi graphs, RGGs and OCNs but not in regular graphs (as their nodes all have the same degree). Thirdly, we focused on the influence of the spatial structure of perturbations by degrading/restoring patches that were either clustered or dispersed in space (Fig. 6). In regular graphs, this was done by drawing the 5 perturbed nodes either from a 9 patch neighborhood (3×3 patches in grid landscapes or 9 adjacent patches in circular landscapes) for the “clustered” modality or drawing them from the whole landscape for the “dispersed” modality. In RGGs and OCN, we identified modules using the function *edge.betweenness.community* (R-package igraph version 1.2.6) and the dispersal matrix *M*, and drew 5 patches from a single modules for the “clustered” modality or from 5 different modules for the “dispersed” modality. We did not conduct this third analysis on Erdös-Rényi graphs as they are not spatially explicit so there is no notion of proximity between patches.

**Figure 2:**
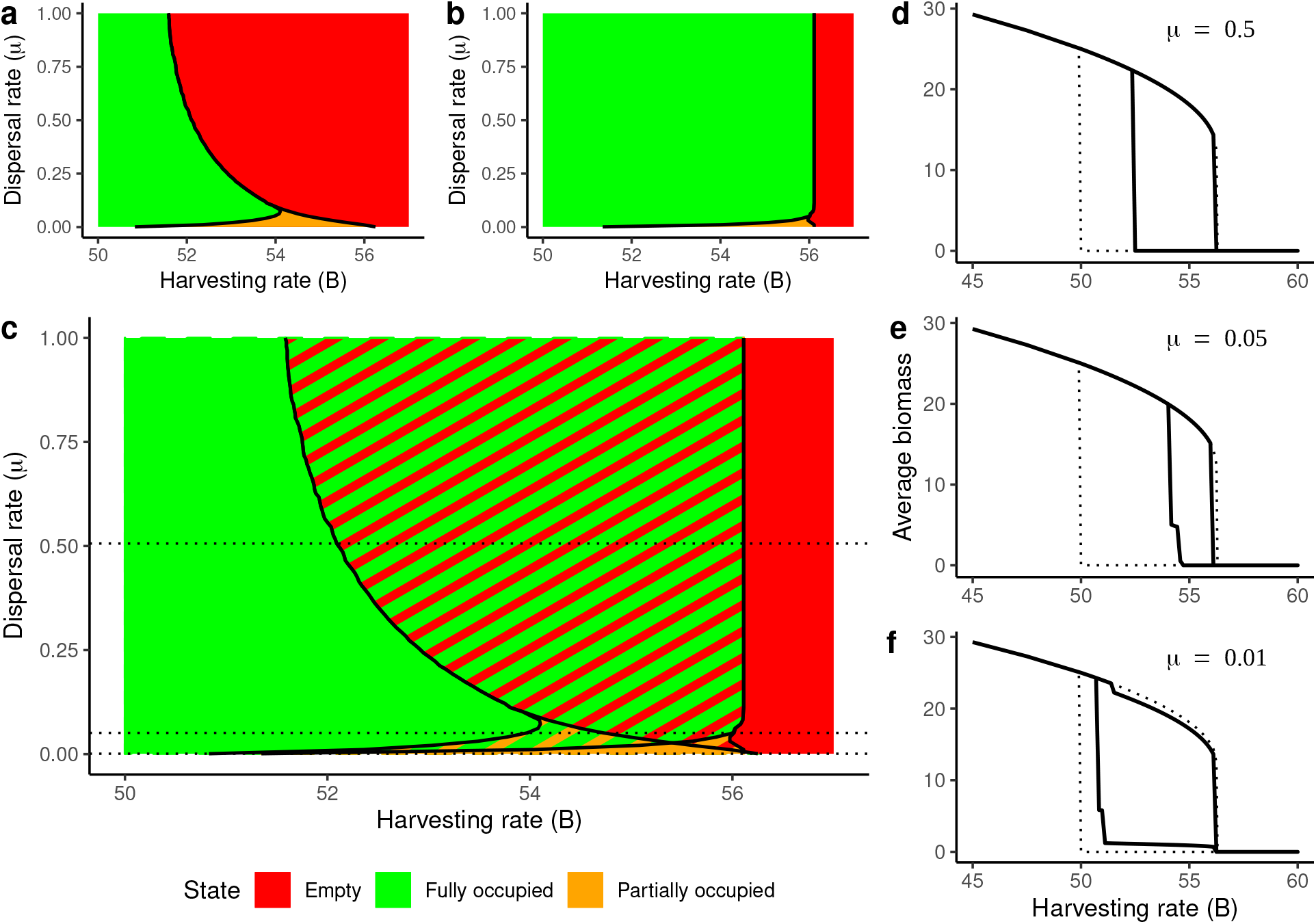
State diagram of a regular grid made of 10*10 bistable patches as a function of the harvesting rate (*B*) and the dispersal rate (*mu*) (a, b, c) and bifurcation diagrams of the average biomass for various dispersal rates (d, e, f). **(a) and (b)** state diagrams of a regular grid of 10*10 bistable patches when starting from 5 random occupied patches in a otherwise empty landscape (a) or 5 random empty patches in an otherwise occupied landscape (b). The colors denote the final state of the system: “fully occupied” (green) means that the total equilibrium biomass was greater than 99% of the maximal equilibrium biomass, “empty” means that the total equilibrium biomass was smaller than 1% of the maximal equilibrium biomass, and “partially occupied” means that the system was neither empty nor fully occupied. **(c)** general state diagram of the aforementioned system, obtained by superimposing panels (a) and (b). **(d), (e) and (f)** bifurcation diagrams of the average biomass of a regular grid (10*10 bistable patches) for different dispersal rates ((d): *μ* = 0.5, (e): *μ* = 0.05, (f): *μ* = 0.001). State space limits from panels (a), (b) and (c) were obtained as the average of 50 replicates; panels (d), (e) and (f) are the bifurcation diagram of a single replicate each.

**Figure 3:**
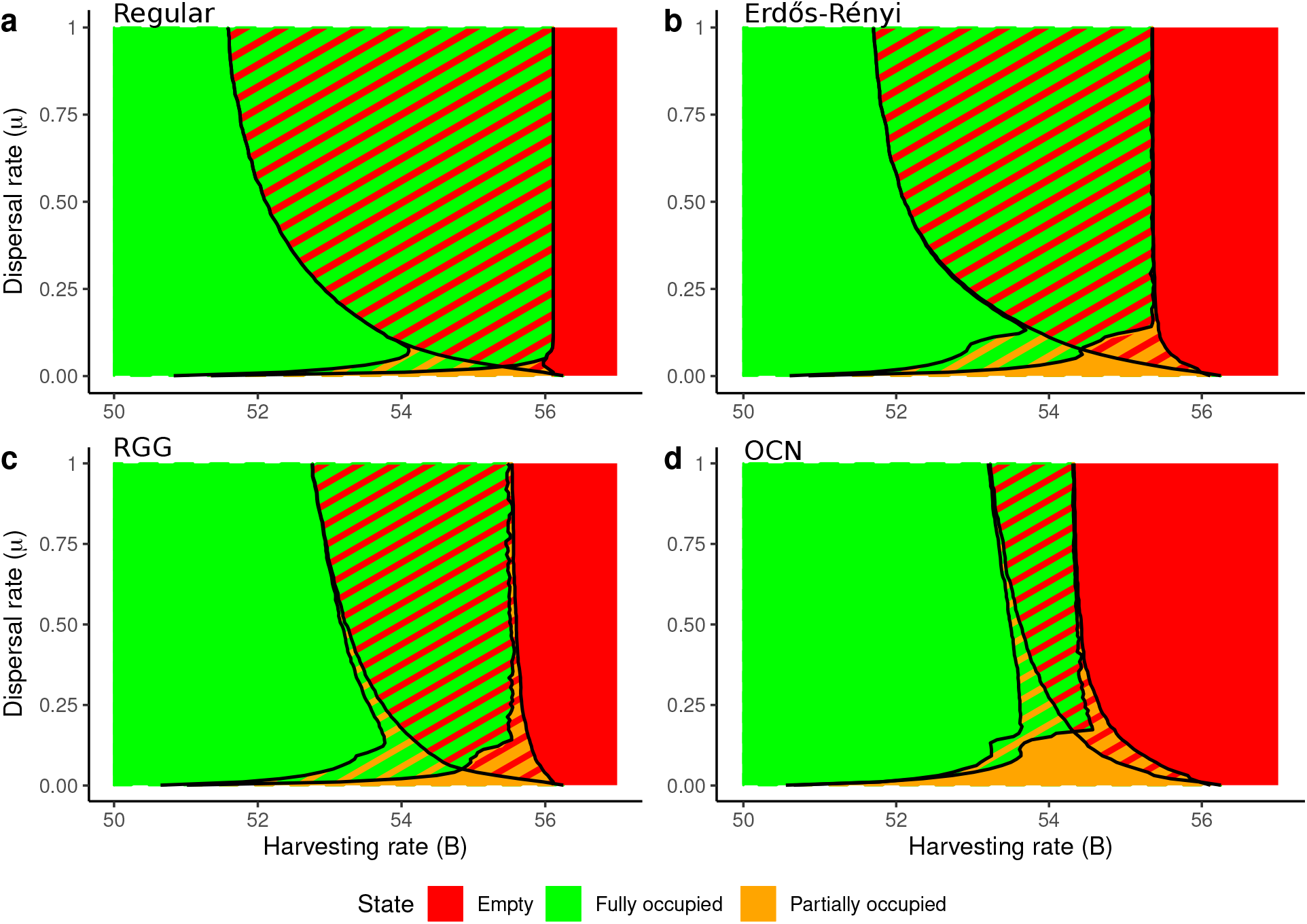
State space diagrams of spatially structured metapopulations as a function of the harvesting rate (*B*) and the dispersal rate (*μ*) for various types of landscapes: (a) a regular grids (10*10 patches on a torus) (same as fig 2c); (b) Erdös-Rényi graphs; (c) random geometric graphs (RGGs) and (d) optimal chanel networks (OCNs). Each panel was obtained by combining the state space diagram of a landscape starting with 5% of random occupied patches and the state space diagram of a landscape starting with 5% of random empty patches, averaged over 50 replicates, similarly as Fig. 2(a-c).

**Figure 4:**
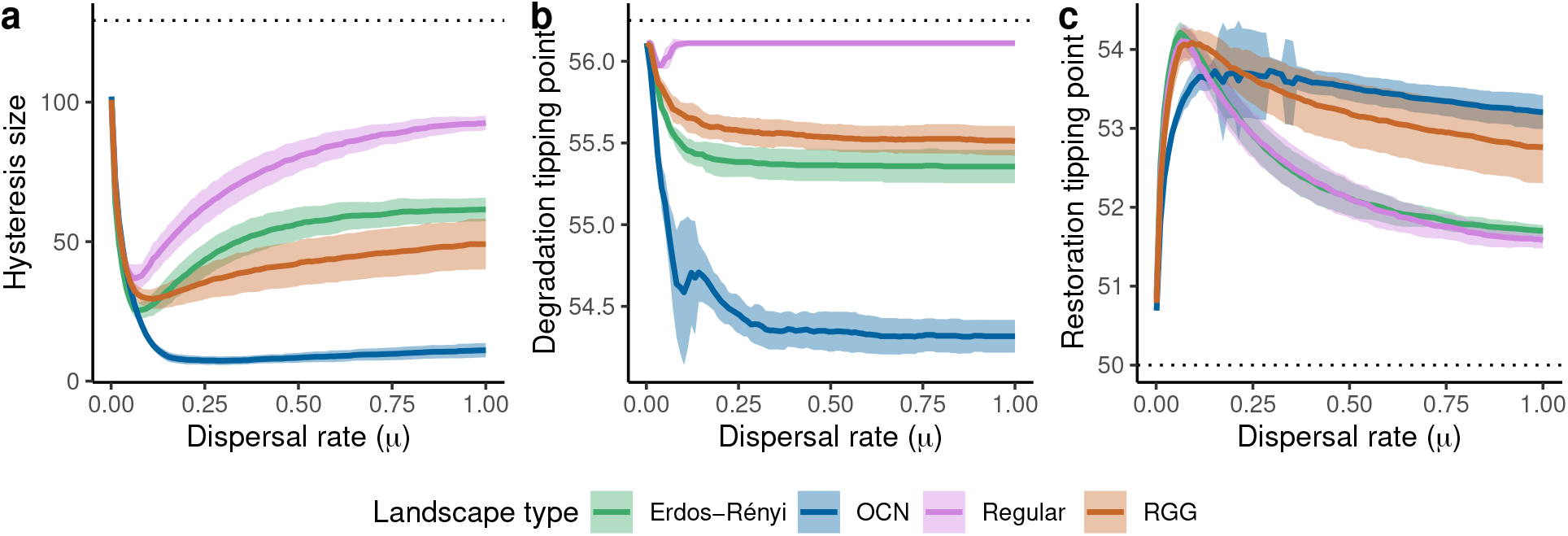
Effect of different landscape types (spatial structures) on the regional dynamics of metapopulations (purple: regular graphs; green: Erdös-Rényi graphs; brown: random geometric graphs (RGGs); blue: optimal channel network (OCNs)). (a) Size of the hysteresis (computed as the area between the upper and lower branches of the bifurcation diagram of the average biomass) as a function of the dispersal rate. (b) and (c) large scale tipping points as a function of the dispersal rate: value of the harvesting rate (*B*) at which the system loses (b) or gains (c) the most biomass. All quantities are computed as the mean (full line) and standard deviation (colored areas) of 50 replicates.

**Figure 5:**
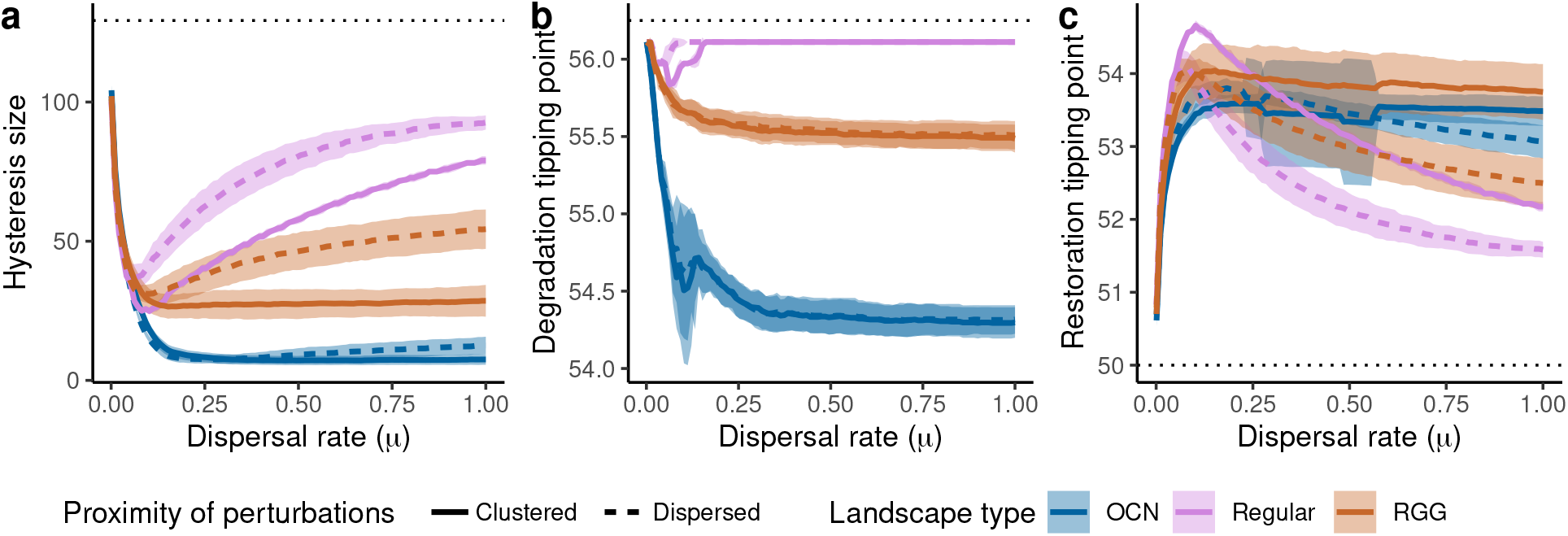
Effects of different perturbation types on the regional dynamics of metapopulations (full line: clustered perturbations, dashed lines: dispersed perturbations) and different landscape types (purple: regular grid; brown: random geometric graphs (RGGs); blue: optimal channel networks (OCNs)). (a) Size of the hysteresis as a function of the dispersal rate. (b) and (c) regional scale tipping points as a function of the dispersal rate: value of the harvesting rate (*B*) at which the system loses (b) or gains (c) the most biomass. All quantities are computed as the mean (lines) and standard deviation (colored areas) of 50 replicates.

**Figure 6:**
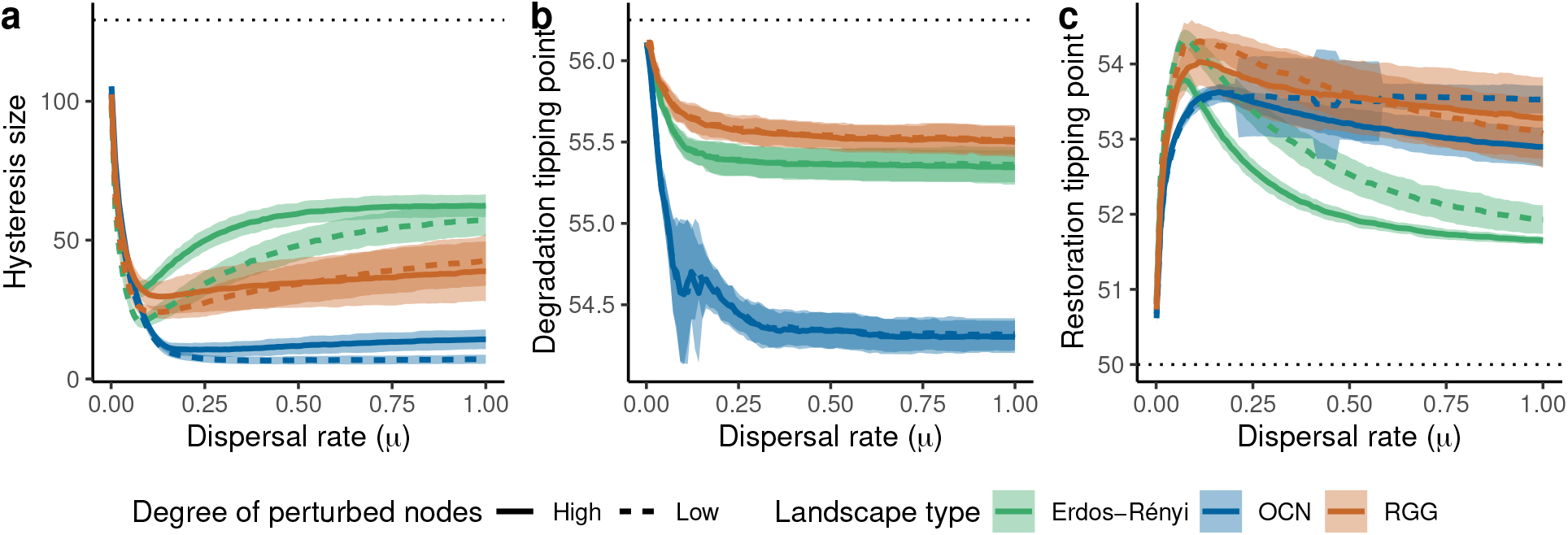
Effect of different perturbation types on the regional dynamics of the metapopulation (full line: perturbation of high-degree patches, dashed lines: perturbation of low-degree patches) and different landscape types (green: Erdös-Rényi graphs; brown: random geometric graphs (RGG); blue: optimal channel networks (OCN)). (a) Size of the hysteresis as a function of the dispersal rate. (b) and (c) regional scale tipping points as a function of the dispersal rate: value of the harvesting rate (*B*) at which the system loses (b) or gains (c) the most biomass. All quantities are computed as the mean (lines) and standard deviation (colored areas) of 50 replicates.

### Characterizing large scale shifts

From the established bifurcation diagrams, we extracted synthetic information on the large scale behaviour of the system.

#### State diagram

We affected one of three categorical states to each simulation depending on its steady-state average biomass: it was considered “fully occupied” when the average biomass was higher than 99% of the pre-perturbation high-biomass equilibrium, “empty” when the average biomass was less than 1% of it and “partially occupied” in between. We constructed state diagrams for each type of networks by averaging the states borders over all replicates and plotting them as a function of the harvesting rate (*B*) and dispersal rate (*μ*).

#### Hysteresis size

We computed the size of the hysteresis as the area in between the higher and lower branches of the bifurcations diagram.

#### Tipping point

Finally, we computed the position of the tipping points from the bifurcation diagrams as the value of harvesting rate (*B*) at which the average biomass decreased (for the degradation trajectory) or increased (for the restoration trajectory) the most.

## Results

### Analysis of a two-patch system

Starting from an isolated patch exhibiting bistability (Fig. 1a) and connecting it to another patch, we found that, at high dispersal, spatial bistability — the stable coexistence of patches in different states — is not possible and the whole system behaves as a single bistable unit (Fig. 1b, supplement S5). As dispersal decreases, spatial bistability becomes possible as new stable states appear (Fig. 1c, supplement S5) with strong differences in biomass between the two patches.

### Stable states of larger metapopulations

We now focus on larger metapopulations made of 100 patches. The results described in the following sections held qualitatively when considering graphs that were overall more or less connected (see sensitivity analysis in supplement S4).

On a regular grid, a larger metapopulation can reach different states (fully occupied, empty or partially occupied) depending on dispersal (*μ*) and harvesting (*B*) rates (Fig. 2). At high dispersal rates (Fig. 2a-d, *μ* > 0.15), only two stable states are possible: the landscape is either fully occupied (the average biomass is that of a single patch) or empty (the average biomass is null), and the metapopulation is bistable as a whole (regional bistability). The bifurcation diagram of the average biomass is then qualitatively similar to that of a single patch (Fig. 2d), albeit with a displaced restoration tipping point. As dispersal rate decreases, the bistability domain shrinks and partially occupied states (where some of the landscape patches are empty and others are occupied) become stable (*i.e*., spatial bistability; Fig. 2c, e). At very low dispersal rates, the system behaves roughly as a collection of independent patches: spatial bistability is common, and local degradation (resp. restoration) can neither spread in space nor be reversed. As a consequence, the bifurcation diagram of the average biomass mostly reflects the initial conditions (Fig. 2f).

These results hold qualitatively for other types of landscapes such as Erdös-Rényi and Random Geometric Graphs (RGG), even though spatial bistability is then more common than for regular grids (Fig. 3a-c). In Optimal Channel Networks (OCNs), we observed a similar state space diagram, although the bistability domain was reduced and did not expand much with increasing dispersal rates, and the partially occupied (*i.e*., spatially bistable) state was stable at higher dispersal rates than for the other networks (up to *μ* ≈ 0.25) (Fig. 3d).

### Characterisation of regional scale catastrophic shifts

#### Hysteresis size

For all landscape types, the hysteresis size of the metapopulation was smaller than that of an isolated patch (Fig. 4a). Hysteresis sizes were similar across network types at low dispersal rates (*μ* < 0.1) but quickly diverged for higher dispersal rates: regular graphs had the larger hysteresis size, followed by Erdös-Rényi graphs and RGGs and finally by OCNs, whose hysteresis 202 size was almost null at high dispersal rates (*μ* > 0.2). For regular graphs, Erdös-Rényi networks 203 and RGGs, we found a unimodal relationship between the hysteresis size and dispersal rate: the 204 hysteresis size quickly decreased at first and reached a minimum at *μ* ≈ 0.1 before slowly increasing with the dispersal rate to reach a plateau (*μ* > 0.1). Hysteresis size in OCNs also decreased with 206 dispersal rate at first, but stayed at a constant value for higher dispersal rates (*μ* > 0.2).

#### Tipping points

For all landscape types, the degradation (resp. restoration) tipping points happened 208 at a lower (resp. higher) harvesting rate than in an isolated patch, which explains the lower hysteresis 209 size in spatially structured landscapes (Fig. 4b and c). The degradation tipping point was mostly 210 unaffected by dispersal rate beyond a threshold (*μ* > 0.1 for regular graphs, Erdös-Rényi graphs 211 and RGGs and *μ* > 0.25 for OCNs). The restoration tipping point, however, was more affected by 212 dispersal rate (Fig. 4c). Interestingly, this means that the patterns observed for hysteresis size are mainly explained by modifications of the restoration rather than of the degradation point. OCNs were the landscapes whose tipping points were overall the most different from an isolated patch.

Similarly to what was observed for hysteresis size, the tipping points were closer to those of a single patch at low dispersal rates, but then deviated from it before reaching a plateau.

### Perturbation modality and large scale catastrophic shifts

Lastly, we considered how different types of perturbations affected large scale shifts, as opposed to the random perturbations considered up to this point.

#### Spatial proximity of perturbations

Both in regular graphs and RGGs, spatially clustered perturbations resulted in a smaller hysteresis compared to spatially dispersed perturbations (Fig. 5a) for all dispersal rates higher than ~ 0.1. This was once again explained by differences in the restoration tipping point: while degradation happened at the same harvesting rate in both perturbation modalities (Fig. 5b), the restoration happened at higher harvesting rates for clustered perturbations (Fig. 5c).

In OCNs, the spatial organization of perturbations had no effect on the hysteresis size (Fig.5a) as both degradation and restoration were mainly unaffected by the perturbation modality (Fig. 5b and c).

#### Connectivity of perturbed patches

In RGG, perturbing highly vs. lowly connected patches had no discernible effect on the hysteresis size and tipping points of the metapopulation (Fig. 6). In Erdös-Rényi graphs and OCNs, perturbing lowly connected patches resulted in a slightly smaller hysteresis at high dispersal rates (*μ* > 0.1) (Fig. 6a), once again because of differences in the restoration point: restoration happened at higher harvesting rate for perturbations of lowly connected nodes compared to highly connected nodes (Fig. 6c), while the degradation point was unaffected (Fig. 6b).

## Discussion

### Stable states of metapopulations composed of locally bistable patches

We found that the stable states of metapopulations made of locally bistable patches depended strongly on dispersal rate and were qualitatively similar for various spatial structures and network connectivity.

Strong dispersal homogenized biomass over space and drove the system towards one of two stable states: either all the patches were occupied and the average biomass was roughly the same as the positive equilibrium of an isolated patch, or all patches were empty and the average biomass was null. These two states had overlapping stability ranges, making the metapopulation bistable as a whole, with a clear hysteresis and abrupt shifts. Therefore, we expect the local degradation or restoration of a few patches in an otherwise homogeneous metapopulation to be quickly reversed, which means that, in a metapopulation with strong dispersal, local restoration efforts are bound to fail unless the environmental conditions cross a metapopulation-scale tipping point. Once this tipping point is crossed, local restoration efforts should spread in space until the whole metapopulation is restored.

When dispersal was sufficiently weak, patches were functionally independent from each other. The exchange of biomass was too low for patches to impact the state of their neighbours and spatial bistability was common as occupied and empty patches could thus coexist in space. In our setting, perturbing only 5% of the patches, this resulted in a large scale behaviour very similar to that of a single patch: the unperturbed patches (95% of the landscape) shifted only when the tipping point of an isolated patch was reached, independently of the perturbations. From a conservation standpoint, the effective independence of patches ensures that a local degradation won’t spread in space and policies could focus on local measures of protection without worrying about spatial effects. However, in the absence of significant fluxes between patches, spatial heterogeneity and stochasticity should have a strong impact on local dynamics and extinction probability. How this can affect large scale dynamics is still an open question as these processes can synergize in surprising ways: for example, Martín et al. (2015) found that demographic stochasticity coupled with a low dispersal rate results in smooth transition in 1- and 2-dimensional continuous space.

Interestingly, this means that in metapopulations made of locally bistable patches, strong dispersal should raise concerns as it translates into possible regional scale catastrophic shifts with a pronounced hysteresis and little prospect for local restoration. On the other hand, weak dispersal should open the door to local conservation and restoration efforts. This should be taken into account when planning conservation measures such as assisted migration or ecological corridors, as increasing a species dispersal rate could make it prone to large scale catastrophic shifts.

### Characteristics of regional scale catastrophic shifts

We observed a smaller hysteresis size than expected from the dynamics of isolated patches for all landscape types. This is particularly pronounced in OCNs where hysteresis was largely reduced (less than 1 / 10^*th*^ of the hysteresis of a single patch for dispersal rates > 0.2). This was expected from previous studies, which predicted that hysteresis should mostly disappear at high dispersal in one dimensional continuous space (van de Leemput et al., 2015) or in linear metapopulations (Keitt et al., 2001; Hilt et al., 2011). Since OCNs are made of large linear parts, they fit this prediction best. The other types of landscapes have more links per patch on average than OCNs, which diluted the effect of any focal patch on more than two neighbours. As a consequence, a local degradation or restoration in an otherwise homogeneous landscape is less likely to spread to its neighbours and more likely to be reversed than in less connected landscapes, resulting in a larger hysteresis.

Interestingly, this reduced hysteresis compared to an isolated patch is mostly due to an earlier restoration, *i.e*., restoration was reached at harsher conditions than expected from the local dynamics. This is due to an asymmetry in the bifurcation diagram of local dynamics: the tipping point towards restoration is a transcritical bifurcation (Fig. 1a, point F1), while the tipping point towards degradation is a saddle-node bifurcation (Fig. 1a, point F2). As a consequence, the null biomass equilibrium has a small basin of attraction while the positive equilibrium has a large basin of attraction, even close to its tipping point, so local restorations are more easily spread in space than local degradations. This asymmetry of the basins of attraction holds true for various models of desertification (Rietkerk and van de Koppel, 1997; Klausmeier, 1999); we therefore expect these systems to also exhibit a reduced hysteresis and easier restoration in metapopulations than expected from the dynamics of a single patch. However, it is important to keep in mind that other models of bistability should behave differently. For example, a system with two transcritical bifurcations should show almost no hysteresis, while a system with saddle-node bifurcations should show an hysteresis of similar size to an isolated patch. Interestingly, a system with a transcritical bifurcation for the degradation and a saddle-node bifurcation for the restoration should have an hysteresis size similar to what we find here, but mainly through an earlier degradation.

Hysteresis size was found to vary with dispersal rate. The largest hysteresis size was observed at the lowest dispersal rate (*μ* = 0.001) for all types of networks, and quickly decreased with dispersal rates. After this initial decrease, the hysteresis size increased with dispersal to a plateau. This means that the smallest hysteresis size and the easiest restoration happened for intermediate dispersal rates (*μ* ≈ 0.1) for which the dispersal is strong enough for a local restoration to affect neighbouring patches, but still sufficiently weak to avoid the dilution of the added biomass in space, triggering a travelling wave of restoration. Interestingly, we observed a large hysteresis size at both very low and high dispersal rates, but with opposite underlying mechanisms: in the case of low dispersal, large scale shifts are hard to induce because local perturbations in a few patches won’t affect the neighbouring patches at all. At high dispersal, large scale shifts are also hard to induce, but because local perturbations quickly get diluted across the whole landscape.

It is worth noting that these variations with dispersal rate are reminiscent of Zelnik et al. (2019)’s regimes of spatial recovery: local restoration efforts could only spread in space at intermediate dispersal rates (what Zelnik called “rescue recovery”), but not at very low dispersal (“isolated recovery”: local restoration does not affect neighbouring patches) or at very high dispersal (“mixing recovery”: the local restoration effort is diluted over the whole landscape).

Characterizing how hysteresis size varies with dispersal rate is important from a conservation standpoint, because a smaller hysteresis means that it is easier to restore the system after a large scale degradation. We can try to derive broad predictions about which species should raise concern for large scale catastrophic shifts. In terrestrial systems, dispersal increases with height for plants (Tamme et al., 2014) and often with body size for animals (Stevens et al., 2014), while population growth rate is expected to decrease with body mass (Savage et al., 2004). Because of these relationships, we can speculate that one should be more concerned by the possibility of large scale catastrophic shifts in systems dominated by large species. Similarly, oceanic systems may be more prone to large scale catastrophic shifts with a marked hysteresis since oceanic species can exhibit strong propagule dispersal that impacts local dynamics (Kinlan and Gaines, 2003). Of course, dispersal also depends strongly on other species traits (*e.g*., seed dispersal syndromes, body temperature) and such broad predictions remain speculations and cannot replace the detailed knowledge on a species of interest.

The landscape type also affected the metapopulation regional scale behaviour. In OCNs, hysteresis size did not increase with dispersal rates but stayed low after the initial decrease. This is because OCNs are almost one-dimensional, with large linear parts. This linearity limited the aforementioned dilution of local restoration efforts and resulted in tipping points almost unchanged by dispersal at high dispersal. On top of that, patches in OCNs have a low average degree, with most nodes having only 1 or 2 neighbours. Because of this, the dispersal from any focal patch was concentrated on a few neighbours. This allowed an easier induction of regional scale catastrophic shifts through a domino effect (even for degradation), as evidenced by the degradation and restoration tipping points happening at respectively lower and higher harvesting intensity than in the other landscape types. Interestingly, two other studies support this result: Hilt et al. (2011) modeled a river as a strictly linear chain of discrete patches and found that hysteresis disappears entirely at high dispersal rates, while Martín et al. (2015) showed that one dimensional systems tend to have smaller hysteresis and smoother transitions than their two (and even three) dimensional counterparts. The other landscape types (grid and RGGs) have no linear parts and an higher average degree (4 neighbours on average). Because of this, the effect of local perturbations was split between several neighbours and diluted in the metapopulation, making it more difficult to induce regional scale shifts and resulting in a larger hysteresis. This was supported by our sensitivity analysis on network connectivity (supplement S4): networks with a smaller average degree (Fig. S4.1) had a smaller hysteresis while networks with a larger average degree (Fig. S4.2) had a larger hysteresis. From these results, we expect large-scale catastrophic shifts with a pronounced hysteresis to be more likely in well-connected and two dimensional systems (*e.g*., terrestrial systems, open ocean) than in systems that are lowly connected or restricted to large linear stretches (*e.g*., rivers, coastal ecosystems).

Lastly, we explored how the spatial structure of perturbations themselves could affect large scale catastrophic shifts. We found that close-by restoration efforts, *e.g*., patches from a single module in RGGs, are more likely to induce regional scale restorations and results in a smaller hysteresis size than restoration efforts conducted randomly in space. This is because restoring close-by patches concentrates the added biomass on a smaller area (*e.g*., the module where we restore patches in RGGs) and limits the dilution of biomass in space. This makes it easier to trigger the large scale restoration of the metapopulation. Studies in continuous space highlight a similar result: van de Leemput et al. (2015) identifies a “minimal size of disturbance to initiate a travelling wave”. Several disturbances smaller than this size won’t initiate a large scale shift as they get diluted individually, but grouping these disturbances over a single area will initiate a large scale shift.

On the other hand, the connectivity of perturbed patches slightly increased the hysteresis size in Erdös-Rényi graphs and OCNs, and had surprisingly no effect in RGGs. This is probably because sampling from low degree patches for the perturbations incidentally results in sampling far away patches, which partially masks the effect of patch connectivity on hysteresis.

### Conclusion

In conclusion, we showed that one can not extrapolate the bifurcation diagram of a single patch to predict regional scale dynamics. Instead, we highlight that a metapopulation can exhibit large scale catastrophic shifts and hysteresis, and that the position of its tipping points depends on both its spatial architecture and its dispersal rate. We find that restoration after a large scale degradation is easier with *i*) local restoration efforts concentrated on a restricted area and *ii*) intermediate dispersal rates. Our findings differ markedly from the predictions established on regular one-dimensional systems and show that we should consider the explicit structure of metapopulations or at least their properties (dimension, average degree and degree heterogeneity) when trying to predict their regional scale dynamics.

## Author contributions

C.S., E.A.F., B.P. and S.K. conceived the study. C.S. and B.P. wrote the model and conducted the simulations. C.S. conducted the mathematical analysis of a two patch system. C.S., E.A.F. and S.K. wrote the manuscript and all authors commented on the draft.

## Data availability

Code is available on GitHub via Zenodo: https://doi.org/10.5281/zenodo.5705432

## Supplementary material

### S1 Metapopulation model

We used a model of spatially structured population describing the dynamics of *n* patches linked by the dispersal of individuals. The dynamics of the biomass (*x_i_*) inside the patch *i* are described by the following differential equation:

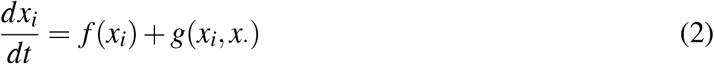

where *f*(*x_i_*) is a function describing the local dynamics of the patch and *g*(*x_i_, x*.) describes the net dispersal between the patch *i* and its neighbors.

#### Local dynamics

We describe the local dynamics with a model coined by Noy-Meir (1975) that exhibits bistability and catastrophic shifts between a null-biomass state (*x_i_* = 0) and a positive equilibrium (*x_i_* = *x*\*) (Eq. 3). It describes the dynamics of a biotic resource *x_i_* (*e.g*., the biomass of plants) that grows logistically with a growth rate *r* and a carrying capacity *K*. This resource is harvested/consumed by a fixed consumer with a Holling type II functional response (with *B* the maximal harvesting rate and *A* the resource biomass at which the harvesting is half of *B*):

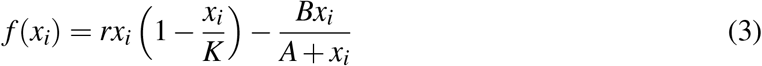

Although it was first coined to describe the dynamics of a plant biomass is a pasture grazed by herbivores (Noy-Meir, 1975), this model is fairly general as logistic growth is commonly used to model the growth of a wide range of organisms (but see Mallet (2012)), and type II functional responses are the norm for numerous consumers (Jeschke et al., 2004; Kalinkat et al., 2013). This system admits two tipping points (*B* = *AR* and *B* = *Ar* + *K*(1 − *AK*)/4). It is bistable in this range (*AR* < *B* < *Ar* + *K*(1 − *AK*)/4) and admits two stable equilibria (*x_i_* = 0 and *x_i_* = *r*_1_) separated by an unstable equilibrium (0 < *r*_2_ < *r*_1_).

#### Spatial dynamics

We consider that the individuals leave the patch *i* at a constant rate *μ* and are equally split between its neighbours. The patch also receives individuals from its neighbours in the same pattern which yields the following equation:

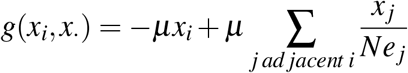

where *Ne _j_* is the number of neighbours of *j*.

#### Global dynamics of the system

The dynamics of the system are thus described by a system of *n* coupled differential equations, which can also be written in a matricial form:

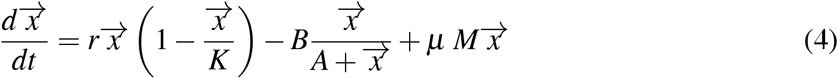

where 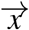 is the vector of local densities and *M* is a matrix describing the structure of the population: the diagonal terms describe the emigration (*M_ii_* = −1) and the off-diagonal terms the immigration (*M_ij,i≠j_* = 0 if *i* and *j* are not neighbors and *M_ij,i≠j_* = 1 /*Ne _j_* if *i* and *j* are neighbors).

#### Two-patch system

The two patch system we used for the analytical stability analysis of a simple spatial system is a particular case of the model presented above, described by the following equations:

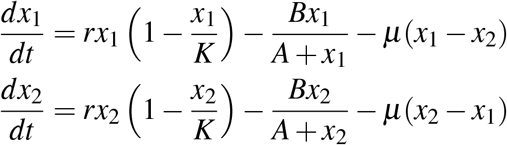

### S2 Landscape design

We generated landscapes of *n* = 100 patches. We used four different landscape structures in order to explore the effect of space on catastrophic shifts: regular graphs (*e.g*., patches arranged either linearly or on a lattice), Erdős–Rényi graphs (where patches are connected randomly), random geometric graphs (RGG, where patches are drawn randomly in space and connected depending on their distance) and optimal channel networks (that are generated using geomorphological processes). The main text analysis is made on networks where the patches have on average 4 neighbors each (when possible), and we conducted sensitivity analysis on the connectivity by generating networks with higher and lower connectivity (section S4).

#### Regular graphs

Regular graphs are graphs in which all nodes have the same number of neighbours: patches are either organized linearly on a cirlce (in 1-D) or on a regular lattice on a torus (for 2-D system). These graph have been used extensively for modelling explicit space (Keitt et al., 2001; van Nes and Scheffer, 2005), so we chose to use them for comparison with existing studies. They can be thought of as a null model of the most ordered spatial system possible. We used patches on a 10*10 lattice with periodic boundary conditions and connections to the 4 nearest neighbours for the main text. For the sensitivity analysis, we used patches on a 10*10 lattice with periodic boundary conditions and connections to the 8 nearest neighbours (2-D system, mean degree = 8) and 100 patches on a circle with connections to the 2 nearest neighbours (1-D system, mean degree = 2).

#### Erdős–Rényi graphs

Erdős–Rényi graphs are a class of graphs obtained by randomly connecting nodes: each pair of patches is connected with a probability *p*. The expected number of links per node (mean degree) is thus *p*∗ (*n* − 1) where *n* is the total number of patches. We used Erdős–Rényi graphs with *n* = 100 patches and a connection probability of *p* = 4/(*n* − 1) for the main text (expected mean degree = 4, realized mean degree = 4). For the sensitivity analysis, we used connection probabilities of *p* = 8/(*n* − 1) (expected mean degree = 8, realized mean degree = 8) and *p* = 2/(*n* − 1) (expected mean degree = 2, realized mean degree = 2.7). After generating a graph, we checked if it was connected (*i.e*., that there existed at least an indirect path between all pairs of nodes) and discarded the graphs that were not connected.

#### Random geometric graphs

Random geometric graphs (RGG) are obtained by drawing nodes randomly in space and connecting the pair of nodes that are closer than chosen a threshold distance. We generated each RGG by drawing 100 pairs of *x* and *y* coordinates uniformly in [0,1]. We then determined the threshold distance that yielded the average degree the closest to a target degree (main text: target degree = 4; sensitivity analysis: target degree = 8). We once again checked if the graphs were connected and discarded those that were not connected. Note that the connectivity of a RGG is constrained by the initial drawing of coordinates, so it was not always possible to reach the targeted average degree. RGGs connectivity deviated a bit from the regular graph counterpart (main text: realized average degree = 4.7; realized degree = 7.9)

#### Optimal channel networks

Optimal channel networks are obtained by simulating geomorphological processes and in order to capture the structural properties of riverine networks. We generated OCN of *n* = 100 patches using the R-package OCNet (version 0.4.0) (Carraro et al., 2020). Because the structure of OCN is constrained by the underlying generating process and generally scale-invariant, we could not manipulate the average degree of these networks (average degree = 1.98).

### S3 Polynomial form of three classical desertification models

In this annex, we highlight the mathematical similarities between four classical models of bistable systems — three models of desertification taken from Noy-Meir (1975), Klausmeier (1999) and van de Koppel et al. (1997); and a model of logistic growth with an Allee effect. We argue that, despite the fact that they are built on different mechanistic processes, they behave similarly as they are all build from a simple third degree polynomial. In the first part, we conduct the exact stability analysis of this third degree polynomial. We then show in the subsequent parts how to relate models of desertification to this polynomial, and how it simplifies their stability analysis.

#### S3.1 Stability analysis of a dynamical model described by a third degree polynomial

The most straightforward way to model bistability is using a cubic polynomial to describe the evolution of a quantity (number of individuals, plant cover, share of users adopting a new technology…) in a closed system. Let us call *P* such a polynomial:

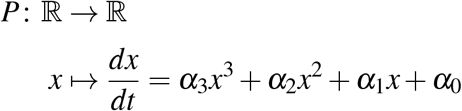

Realistically, *α*_3_ should be negative (otherwise *dx/dt* → ∞ when *x* is high, meaning the population is unbounded). Hence, *α*_3_ is restricted to negative values in what follows. In biological systems, realism also imposes a restriction on *α*_0_: since the system is closed and *x* describes a density of organisms that cannot be spontaneously generated, *P*(0) must be null, and hence *α*_0_ = 0. Additionally, *x* can only be positive when describing a population, so *P* only needs to be defined on 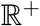.

Hereafter, *P* refers to the previously defined polynomial with these additional restrictions:

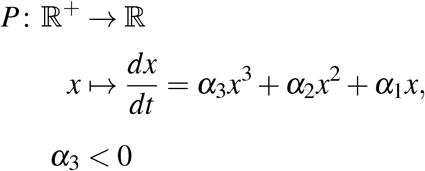

##### Existence of alternative equilibria

Since *α*_0_ = 0, the system has an immediate equilibrium for *x* = 0. The other equilibrium can thus be found by solving:

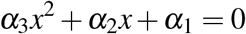

This is a degree two polynomial with discriminant 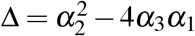. If Δ < 0 (*i.e*. 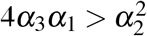), the polynomial has no real root and 0 is the only equilibrium of this system (Fig.S3.1c) If Δ > 0, then there are two other roots to the system (that are only relevant when they are positive):

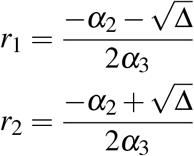

The nature of the system depends on the sign of these roots: if both are negative, then 0 is a stable equilibrium and the only biologically significant equilibrium. If both are positive, 0 is still a stable equilibrium but there is also a stable positive equilibrium (*r*_1_) separated from 0 by an unstable equilibrium (*r*_2_) and thus the system is bistable (Fig. S3.1d). If only one of them is positive, then 0 is unstable and *r*_1_ is stable and the only positive equilibrium (Fig. S3.1e).

##### Sign of *r*_1_

Since *α*_3_ is negative, *r*_1_ is of the same sign as 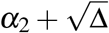. Thus, *r*_1_ is positive iff 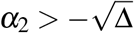

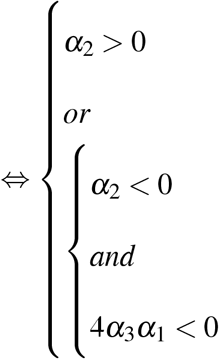

Thus *r*_1_ is only negative when *α*_2_ < 0 and 4*α*_3_*α*_1_ > 0, *i.e*. in the top left corner of the *α*_2_ − 4*α*_3_*α*_1_ plane (Fig.S3.1a).

##### Sign of *r*_2_

Similarly, the sign of *r*_2_ is that of 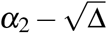. *r*_2_ is positive iff 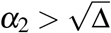

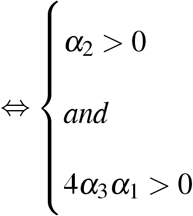

*r*_2_ is thus only positive in the top right corner of the *α*_2_ − 4*α*_3_*α*_1_ plane (FigS3.1a).

These results are summed up in Fig.S3.1:

- if *α*_1_ is positive (crossed area in Fig.S3.1a), the system always reach a stable positive equilibrium (*r*_1_, Fig.S3.1g)
- If *α*_1_ is negative, the behaviour of the system depends on the relative values of *α*_2_ and 4*α*_1_*α*_3_: if 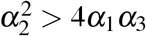 (white area in Fig.S3.1a), the system is bistable and can either reach 0 or a positive equilibrium (*r*_1_) depending on its initial state (Fig.S3.1f). If 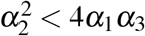, the system has no strictly positive equilibrium and necessarily goes to 0.

##### Stability analysis and interpretation

One way to make light of this is to see that *α*_1_ drives the system behaviour when *x* is close to 0 while *α*_2_ and *α*_3_ become more predominant at high *x* value, thus:

- if *α*_1_ is positive, *P* is positive when *x* is close to 0 and thus 0 is unstable. Since *P* is negative when *x* is high (because *α*_3_ is negative), *P* necessarily becomes null once (and only once) for a *x* > 0 which is a stable equilibrium.
- If *α*_1_ is negative, *P* is negative around 0 and thus 0 is a stable equilibrium. In that case, either *α*_2_ makes a positive contribution high enough (meaning 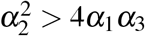) for *P* to reach 0 two more times (because *P* ultimately decreases to –*infinity* at high *x*) and the system is bistable or *α*_2_ contribution is not enough 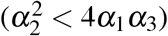 to bring *P* to 0 again and thus 0 is the only equilibrium.

An intuitive way to sum these relationships is by looking at the *α*_2_-4*α*_1_*α*_3_ plane (Fig. S3.1a): in the area under the x-axis (in green), the system necessarily goes to a positive equilibrium. Over the x-axis, the curve defined by *y* = *x*^2^ separates two areas: on the left (in red) where 0 is the only stable equilibrium and the system goes to extinction. On the right (in orange), the system is bistable. By increasing *α*_2_ (moving from left to right) or *α*_1_ (moving from top to bottom), we drive the system away from extinction and towards either a positively stable domain (in green) or a bistable domain (in orange) respectively.

#### S3.2 Application to biological systems

While some theoretical studies chose to use a third degree polynomial as a simple way to obtain bistable dynamics, models rooted in biological mechanisms are rarely expressed as third degree polynomials — because cubic terms cubic terms are hard to intuitively link to a process. Instead the dynamics of biological systems (*dx*/*dt*, usually expressing biomass or species density) are often expressed as a combination of:

- linear terms (*λ* ∗ *x*) to describe density independent processes (*e.g*., death rate in birth-death 602 processes, growth in an exponential model).
- square terms to describe density dependent processes such as mass action law (*e.g*., intraspecific competition in a logistic growth model: *r*(*x* − *x*^2^/*K*)).
- a quotient of low order polynomials (1 or 2) to describe saturating processes (*e.g*., type 2 functional response: *n*/(1 + *n*)).

**Figure S3.1:**
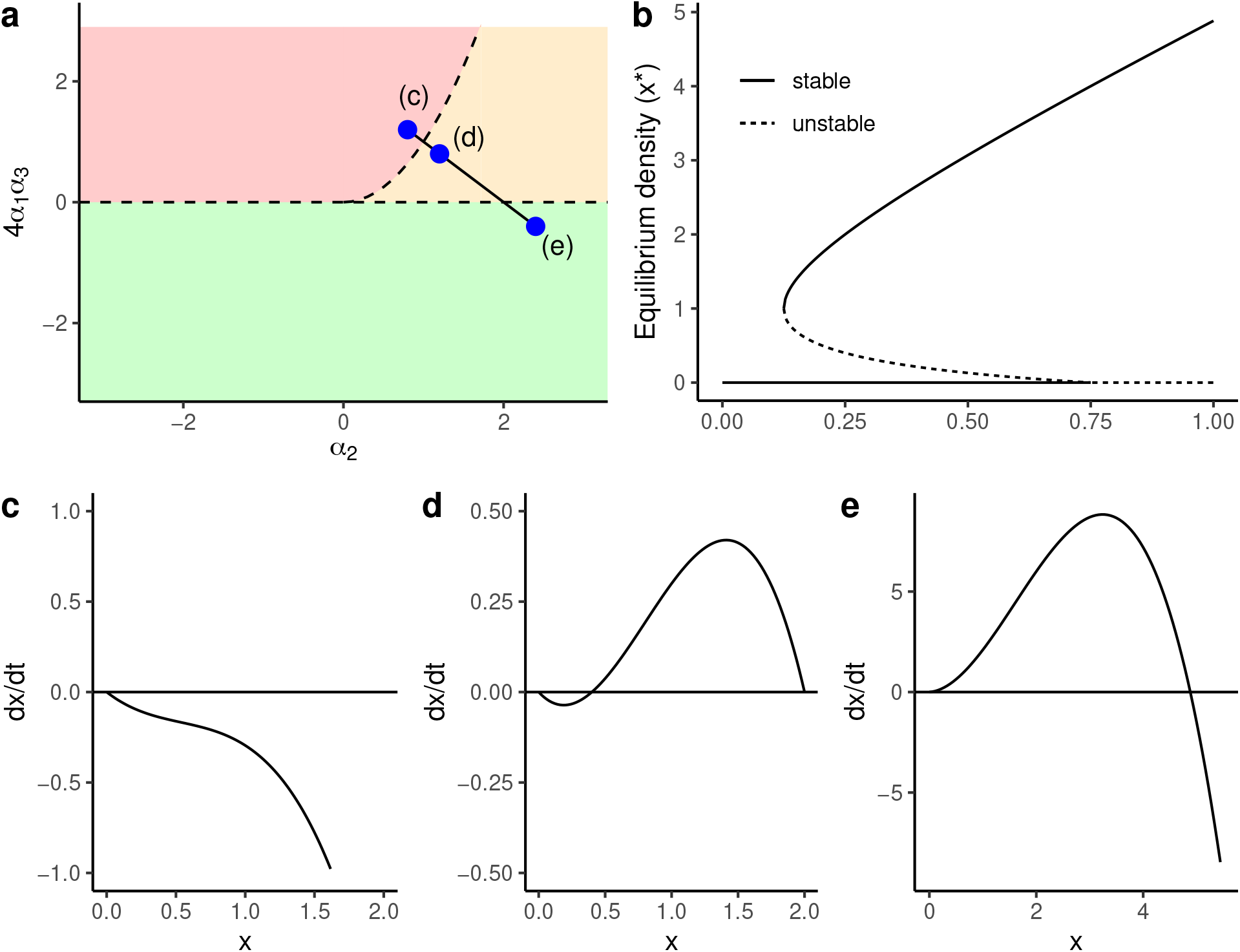
Description of *P* behaviour. **(a)** Phase diagram of *P* depending on *α*_2_ and 4*α*_3_*α*_1_. In the red area, 0 is the only equilibrium in 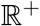 and the system always goes to extinction (see panel c). In the orange area, the system has three equilibrium in 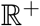 ([0, *r*_1_, *r*_2_]) and goes either to extinction or *r*_2_ depending on the initial *x* (see panel d). In the green area, 0 is an unstable equilibrium and the system always goes to *r*_1_ that is stable (see panel e). The full line shows a transect happening when increasing both *α*_2_ from 0.8 to 2.4 and *α*_1_ from −0.6 to 0.2 (*α*_3_ is fixed at −0.5) and the dots show where the panels c, c and e where taken. **(b)** bifurcation diagram (equilibria of *P*(*x*)) along the transect represented in the panel a (the *x* axis figures the position along the line: 0 is the point (c) and 1 is the point (e)). **(c)** a monostable parameterization of *P*(*x*) resulting in extinction (parameters values of the point (c) in panel a: *α*_1_ = −0.6; *α*_2_ = 0.8; *α*_3_ = −0.5. **(d)** a bistable parameterization of *P*(*x*) (parameters values of the point (d) in panel a: *α*_1_ = −0.4; *α*_2_ = 1.2; *α*_3_ = −0.5. **(e)** a monostable parameterization of *P*(*x*) resulting in a positive equilibrium (parameters values of the point (e) in panel a: *α*_1_ = 0.2; *α*_2_ = 2.4; *α*_3_ = −0.5.

**Figure S3.2:**
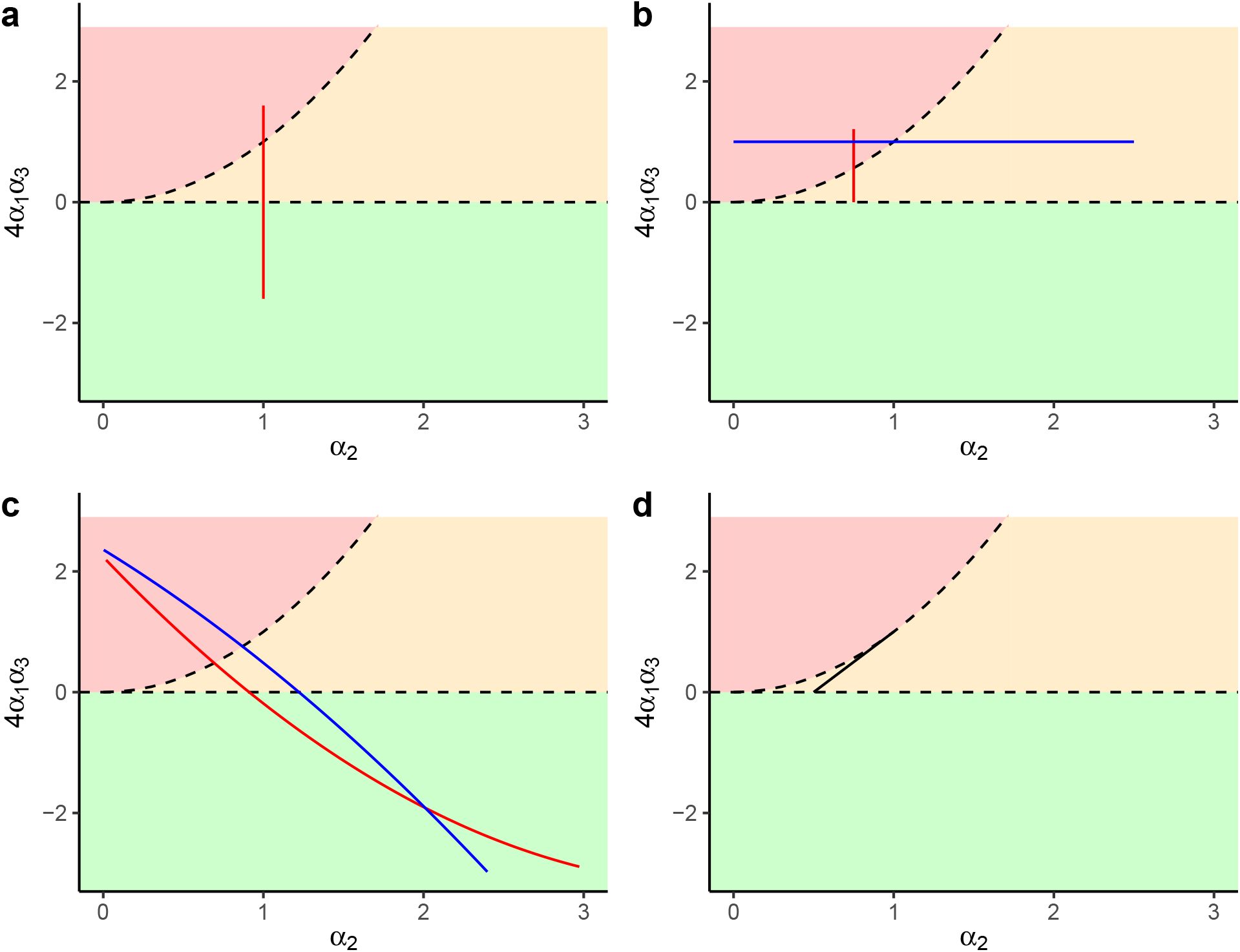
Situation of 4 classical models in the *al pah*_2_-*α*_1_*α*_3_ plane (a: Noy-Meir, b: Klausmeier, c: Rietkerk and Van de Koppel, d: Logistic growth with an Allee effect). Red lines are transect obtained by varying mortality parameters, blue line are obtained by varying water input parameters. **(a)** Effect of varying the grazing pressure (*B* in 40-60) in the Noy-Meir model (other parameters: *r* = 2, *K* = 50, *A* = 25). **(b)** Effect of varying plant mortality (*M* in 0-1.1, red line) or the water input (*A* in 0-10, blue line) in the Klausmeier model (other parameters: *L* = 0.5, *R* = 0.5, *J* = 0.5; *A* = 3 for the red line and *M* = 1 for the blue line). **(c)** Effect of varying plant mortality (*M* in 0-0.1, red line) or the water input (*I* in 4-7, blue line) in the Rietkerk and Van de Koppel model (other parameters: *q* = 0.1, *L* = 1, *U* = 1, *R* = 1, *K* = 10, *A* = 5, *B* = 5; *I* = 4 for the red line and *M* = 0.1 for the blue line). **(d)** Effect of varying the Allee effect threshold (*c* in 0-1) in a logistic model with an Allee effect (other parameters: *r* = 0.5).

We found that a number of these models can be restated as a quotient of two polynomials:

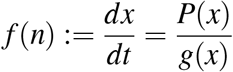

where *P* is the third degree polynomial described earlier and g is a strictly positive and monotonously increasing second order (at most) polynomial (defined on 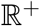). Because of these properties, *P* and *f* share the same sign and roots despite *f* not being a polynomial. In the following section, we illustrate this using three classical models of desertification as well as a model of logistic growth with an Allee effect. We argue that: *i)* the stability analysis of these models is made much easier by mapping *f* to *P* and using the stability analysis of *P*, for which we know the exact expression and nature of its equilibria; *ii)* that expressing the parameters of *P* (*α*_1_, *α*_2_ and *α*_3_) as aggregates of the parameters of *f* helps to understand the effect of biological parameters on the systems equilibria — in particular through the graphical analysis of the *α*_2_-*α*_1_*α*_3_ plane — and *iii)* that the models we consider all share a common structure despite the fact that they describe different mechanistic processes, which is encouraging for the transposability of studies made on one of these models.

##### S3.2.1 Noy-Meir model

The model we use in the main text of this study describes the dynamics of plants (whose biomass is written *x*) grazed by a constant population of consumer (whose maximal grazing rate is *B*), as coined by Noy-Meir (1975). The variations of plant biomass are expressed by the following differential equation:

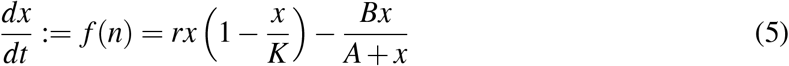

In the absence of grazing, the plant biomass grow logistically with a growth rate *r* up to a carrying capacity *K* (left term). On top of that, a fixed consumer harvest the plant population following a Holling type 2 harvesting rate (right term) where *B* is the maximal harvesting rate and *A* is the biomass of plants at which the per plant biomass harvesting rate is half of its maximal value (half-saturation constant of the grazers): the grazing increases linearly with *x* at low plant biomass then reaches a plateau as each harvesting agent reaches its maximal harvesting rate.

###### Polynomial form

We rewrite *f*(*x*) as a quotient of polynomial:

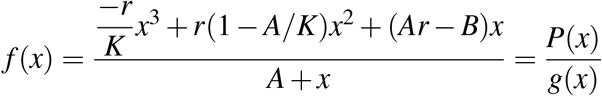

Hence the coefficients of *P*(*x*) are:

**Table S3.1:** Coefficient of the polynomial *P*(*x*) describing the behaviour of the Noy-Meir model.

| Coefficient | Expression |
| --- | --- |
| $\alpha_1$ | $Ar - B$ |
| $\alpha_2$ | $r(1 - A/K)$ |
| $\alpha_3$ | $-r/K$ |

The main control parameter — grazing intensity (*B*) — is only found in *α*_1_. Varying this parameter moves the system vertically in the *α*_2_-4*α*_1_*α*_3_ plane (Fig. S3.2a): it goes from the domain with a single positive equilibrium (green area) to the domain with a single null equilibrium (extinction, red area) through the bistable domain (orange area) as *B* increases. Note that in the unlikely case where *A* > *K* (*i.e*., the grazing saturates at a biomass level greater than the carrying capacity), *α*_2_ is negative so the system goes from a positive biomass to extinction without going through a bistable phase as *B* increases.

##### S3.2.2 Klausmeier model

This model coined by Klausmeier (1999) and describes catastrophic shifts in arid grassland, but here the bistability arises from positive feedbacks between plant biomass (*x*) and water availability (*w*):

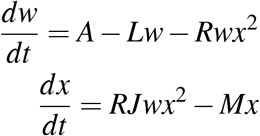

with the following paramters:

- *A* the water input
- *L* the water loss rate
- *R* the rate at which plants uptake water
- *J* a conversion rate from water mass to plant biomass (*i.e*., how much plant biomass is created from 1 unit of water mass).
- *M* the plant mortality rate.

The water dynamics depend on a constant water input (*A*, *e.g*., the precipitation rate), water losses that are proportional to the amount of available water (*Lw*, *e.g*., evaporation and infiltration in deep soil layers) and a water uptake by plants (*Rwx*^2^) that is proportional to both the amount of proportional to the amount of available water and the square of plant biomass (the square term representing the facilitation between plants allows the system to be bistable). The plant dynamics are described by a growth term that is proportional to the water uptake (*RJwx*^2^) and a death rate proportional to plant biomass (*Mx*).

It is common to assume that water dynamics are way faster than plant dynamics in models describing plant-water dynamics. We can thus calculate the water equilibrium w* by solving the equation 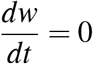:

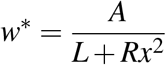

By replacing *w* by its equilibrium *w*\* in the expression of water dynamics, we can describe the system using a single equation:

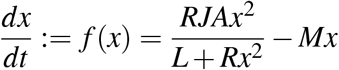

###### Polynomial form

Similarly to the Noy-Meir model, we can rewrite the equation above as a quotient of polynomials:

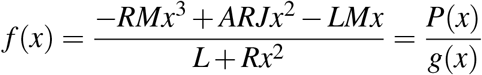

The coefficients of *P*(*x*) are:

**Table S3.2:** Coefficient of the polynomial *P*(*x*) describing the behaviour of the Klausmeier model.

| Coefficient | Expression |
| --- | --- |
| $\alpha_1$ | $-LM$ |
| $\alpha_2$ | $ARJ$ |
| $\alpha_3$ | $-RM$ |

The two main control parameters here are the plant mortality (*M*) and the water input (*A*). The plant mortality is only present in the *al pha*_1_ and *al pha*_3_ so — similarly to the Noy-Meir model — varying *M* moves the system vertically in the *α*_2_-4*α*_1_*α*_3_ plane (Fig. S3.2b): it goes from the frontier between the domain with a single positive equilibrium (green area) and the bistable domain (orange area) to the domain with a single null equilibrium (extinction, red area) as *M* increases. The water input parameter (*A*) is only present in *α*_2_, so varying it moves the system horizontally in the *α*_2_-4*α*_1_*α*_3_ plane (Fig.S3.2b): increasing *A* moves the system from the red domain (monostable, extinction) to the orange domain (bistable).

Note that for this model, *α*_1_ and *α*_3_ are always negative (hence 4*α*_1_*α*_3_ is always positive) and *α*_2_ is always positive. Thus the system is necessarily in the top-right quadrant of the *α*_2_–4*α*_1_*α*_3_ plane and thus is either bistable or admits only the null equilibrium (it cannot reach the green domain in the *α*_2_–4*α*_1_*α*_3_ plane).

##### S3.2.3 Rietkerk & Van de Koppel model

The third model of desertification that we examine was coined van de Koppel et al. (1997) and also describes the dynamics of soil water (*W*) and plant biomass (*x*):

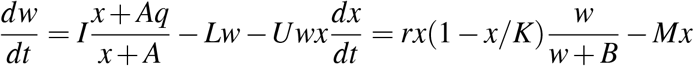

Water (*w*) enters the soil through an infiltration facilitated by plants presence (*I*(*x*+*As*)/(*x*+*A*), with *I* the water input) and exits it through abiotic losses (−*LW*) and plant uptake (−*Uwx*). The plants (*x*) grow following a logistic term (*rx*(1 − *x*)/*K*) multiplied by a saturating function of *w* (*w*/(*w* + *B*)) describing how water affects plant growth; and they lose biomass at a constant rate *M* (−*Mx*). Similarly to the Klausmeier model, we can find the water equilibrium *w*\* by solving *dw/dt* = 0:

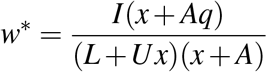

We can inject this equilibrium (under the assumption that water dynamics are faster than plant dynamics) into the expression of dx/dt to find:

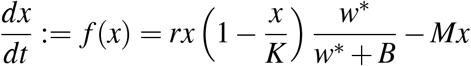

###### Polynomial form

By developing this expression, we find that it is once again a quotient of polynomials:

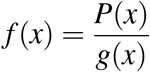

with:

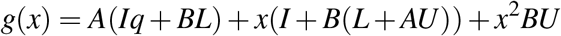

and:

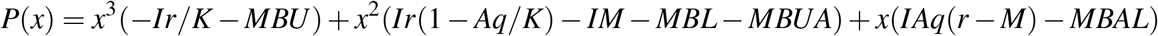

whose coefficients are:

**Table S3.3:**
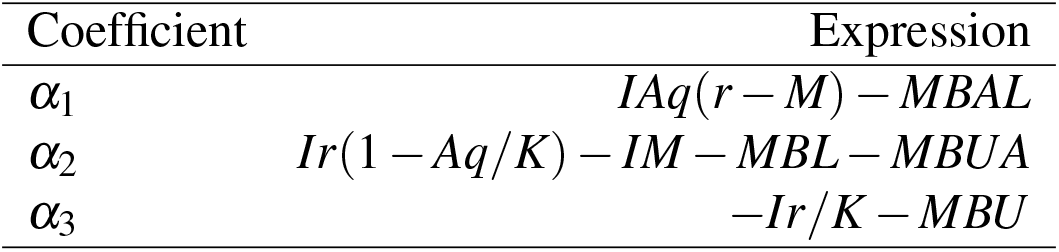
Coefficient of the polynomial *P*(*x*) describing the behaviour of the model described by Rietkerk and Van de Koppel.

This model is a bit more complicated than the previous ones and would warrant a lengthy analysis of all its parameters and how they affect the systems equilibria. However, we will here focus on two simple results:

Firstly, the polynomial form provides a simple way to analyse the system: while the model uses 9 parameters, we can reduce the complexity of its analysis through the three compound parameters *α*_1_, *α*_2_ and *α*_3_. For any set of parameters, it is easy to compute the *alpha* and then *i)* to determine the number and nature of the equilibria through the *α*_2_-4*α*_1_*α*_3_ plane (Fig. S3.2c) and *ii)* to determine the exact values of these equilibrium through the analytical expresions of *r*_1_ and *r*_2_ that we derived in the section S3.1.
Secondly, we can analyze the impact of varying a given parameter using the *α*_2_-4*α*_1_*α*_3_ plane. For the set of parameters used in Fig. S3.2c, we can see that the two control parameters (mortality rate *M* and water input *I*) have a similar effect on the system. The system moves from a stable positive equilibrium (green area) towards extinction (red area) through a bistable state (orange area) when increasing the mortality (*M*, red line in Fig. S3.2c) or decreasing the water input (*I*, blue line).

##### S3.2.4 Logistic growth with an allee effect

Lastly, we focus on the model of a logistic growth with an Allee effect. For simplicity, we use a model where *x* represents the species density rescaled to the carrying capacity (*i.e*., *x* is the species density in proportion of the carrying capacity). This system is described by the following equation:

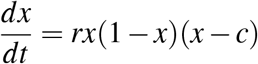

where *r* is the growth rate and *c* is the Allee effect threshold, *i.e*., the density under which the species growth is negative.

###### Polynomial form

Here, the equation describing the system is already a third degree polynomial (*e.g*., g(x) = 1) that we can express by developing the expression above:

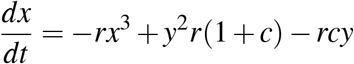

Hence the coefficients of the polynomial form are:

**Table S3.4:** Coefficient of the polynomial *P*(*x*) describing a logistic growth with an Allee effect.

| Coefficient | Expression |
| --- | --- |
| $\alpha_1$ | $-rc$ |
| $\alpha_2$ | $r(1+c)$ |
| $\alpha_3$ | $-r$ |

Because this model is very simple by design, the polynomial analysis is not very useful here. It is however interesting to note that it fits into the same framework as the more mechanistic models presented above. The figures S3.2d shows how this model is situated in the *α*_2_-4*α*_1_*α*_3_ plane as *c* varies.

### S4 Sensitivity analysis on network connectivity

We conducted a sensitivity analysis on network conductivity by conducting the same simulations as in the main text but on networks with higher or lower connectivity (see S2 for the generating methods). We generated regular graphs where each patch had exactly 2 or 8 neighbours, which respectively results in 100 patches on a circle with connections to the 2 nearest neighbours (1-D system, mean degree = 2) and a 10*10 lattice with periodic boundary conditions and connections to the 8 nearest neighbours (2-D system, mean degree = 8). We generated Erdös-Rényi graphs with a target connectivity of 2 or 8, but because we constrained them to be connected graphs, this resulted in an average connectivity slightly higher than 2 (2.7 on average) for the low connectivity networks. We also generated RGGs with a target connectivity of 8 (which resulted in a realized connectivity of 7.9 on average), but couldn’t make RGGs with a lower connectivity than in the main text because of constraints on the generating process.

#### S4.1 General patterns

The general patterns observed in the main text stayed qualitatively unchanged (Fig. S4.1 and S4.2): hysteresis size first decrease with dispersal rate (until *μ* ~ 0.1) and then increased to a plateau.

The effect of landscape type on hysteresis was conserved for highly connected graphs (regular graphs >Erdös-Rényi graphs >RGGs, fig. S4.2) We suspect that this is due to differences in degree distributions: in regular graphs, all patches have the same degree (4 or 8 neighbours), which explains their larger hysteresis. Erdös-Rényi networks and RGGs have some variance in their node degrees, so the perturbations were likely to happen on a low degree node and to initiate a large scale shift, explaining their hysteresis size in between that of regular graphs and of OCNs. The difference between RGGs and Erdös-Rényi graphs could be also due to differences degree distribution: while they have a similar average degree, RGGs have a wider degree distribution, making it more likely that a pertubation happens on a low-degree node than in Erdös-Rényi graphs, hence their smaller hysteresis.

Hysteresis size was similar accross landscapes types for lowly connected graphs (Fig. S4.1), which we conjecture is due to the very low degree heterogeneity at low connectivity.

Lastly, hysteresis size was smaller in less connected networks (Fig. S4.1) and higher in more connected networks (Fig. S4.1), which is consistent with our interpretation that biomass dilution in highly connected networks prevent the spread of local shifts.

#### S4.2 Perturbation modality

Lastly, the effect of perturbation modality was similar to what we observe in the main text. Targeting clustered patches resulted in a smaller hysteresis than targeting patches dispersed over the landscape in both lowly and highly connected networks (Fig. S4.3 and S4.4).

Targeting high or low degree node had only a marginal effect in lowly-connected Erdös-Rényi networks and no effect otherwise (Fig. S4.5 and S4.6), similarly to what we observed in 4-degree networks.

**Figure S4.1:**
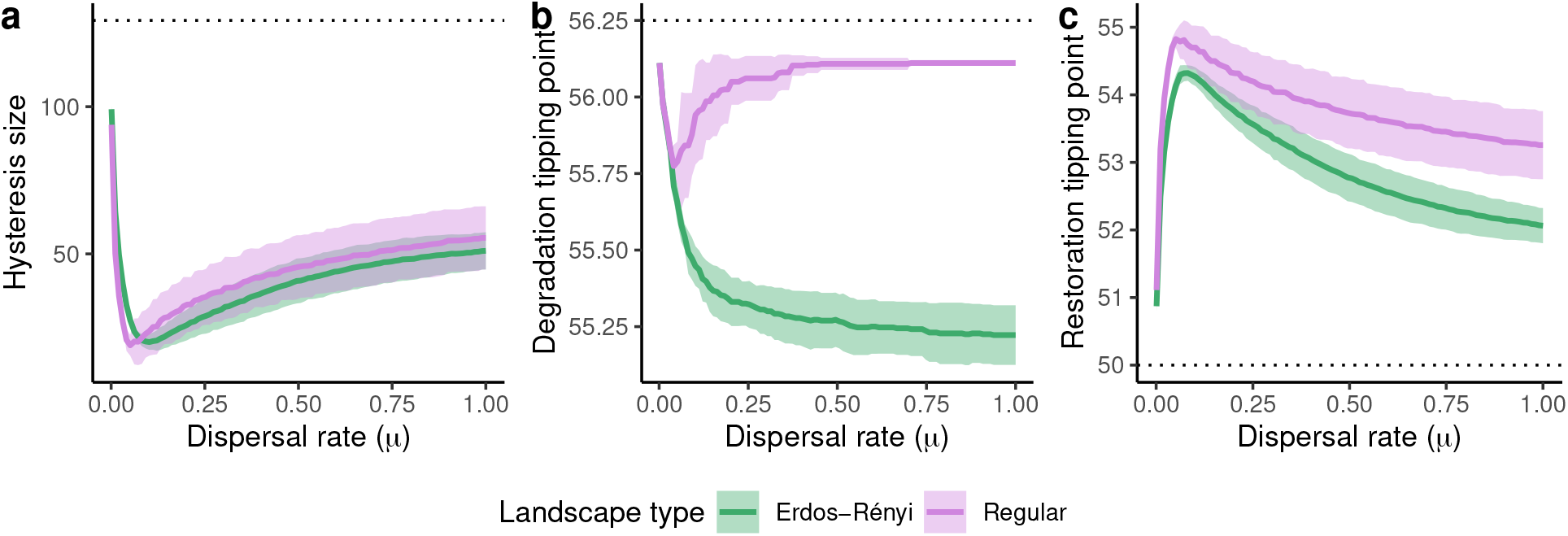
Sensitivity analysis: networks with an average degree of 2. Characterization of the large scale dynamics for different spatial structures of metapopulation (purple: circular network; green: Erdös-Rényi graphs). (a) Size of the hysteresis (computed as the area between the upper and lower branches of the bifurcation diagram of the average biomass) as a function of the dispersal rate. (b) and (c) large scale tipping points as a function of the dispersal rate: value of the harvesting rate (*B*) at which the system loses (b) or gains (c) the most biomass. All quantities are computed as the mean (full line) and standard deviation of 50 replicates.

**Figure S4.2:**
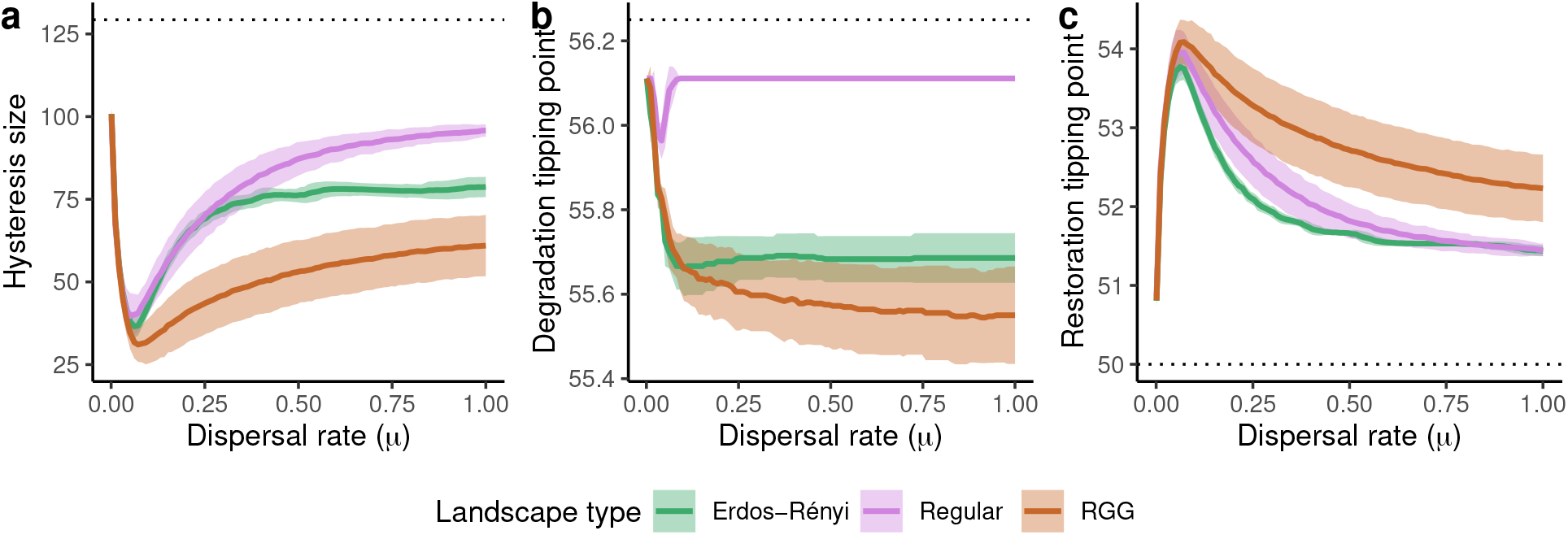
Sensitivity analysis: networks with an average degree of 8. Characterization of the large scale dynamics for different spatial structures of metapopulation (purple: regular grid; green: Erdös-Rényi graphs; brown: random geometric graphs). (a) Size of the hysteresis (computed as the area between the upper and lower branches of the bifurcation diagram of the average biomass) as a function of the dispersal rate. (b) and (c) large scale tipping points as a function of the dispersal rate: value of the harvesting rate (*B*) at which the system loses (b) or gains (c) the most biomass. All quantities are computed as the mean (full line) and standard deviation of 50 replicates.

**Figure S4.3:**
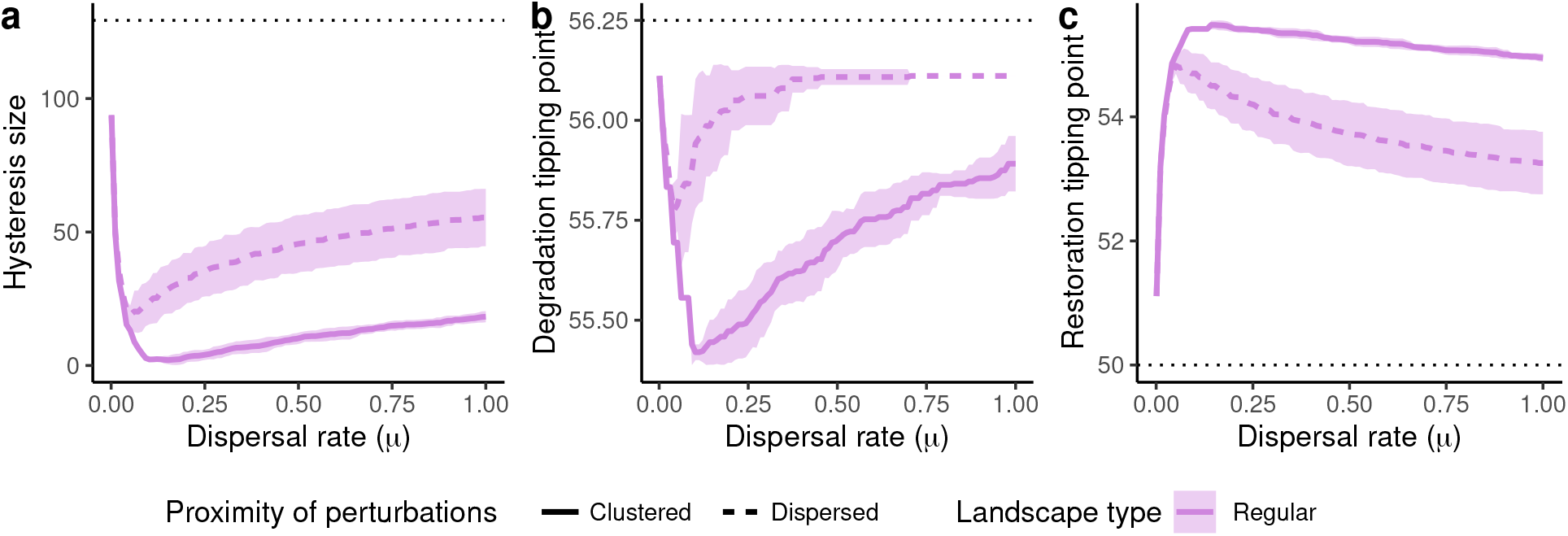
Sensitivity analysis: networks with an average degree of 2. Characterization of the large scale dynamics for different types of perturbations (full line: clustered perturbations, dashed lines: dispersed perturbations) and different spatial structures of metapopulation (purple: circular network). (a) Size of the hysteresis as a function of the dispersal rate. (b) and (c) large scale tipping points as a function of the dispersal rate: value of the harvesting rate (*B*) at which the system loses (b) or gains (c) the most biomass. All quantities are computed as the mean (lines) and standard deviation of 50 replicates.

**Figure S4.4:**
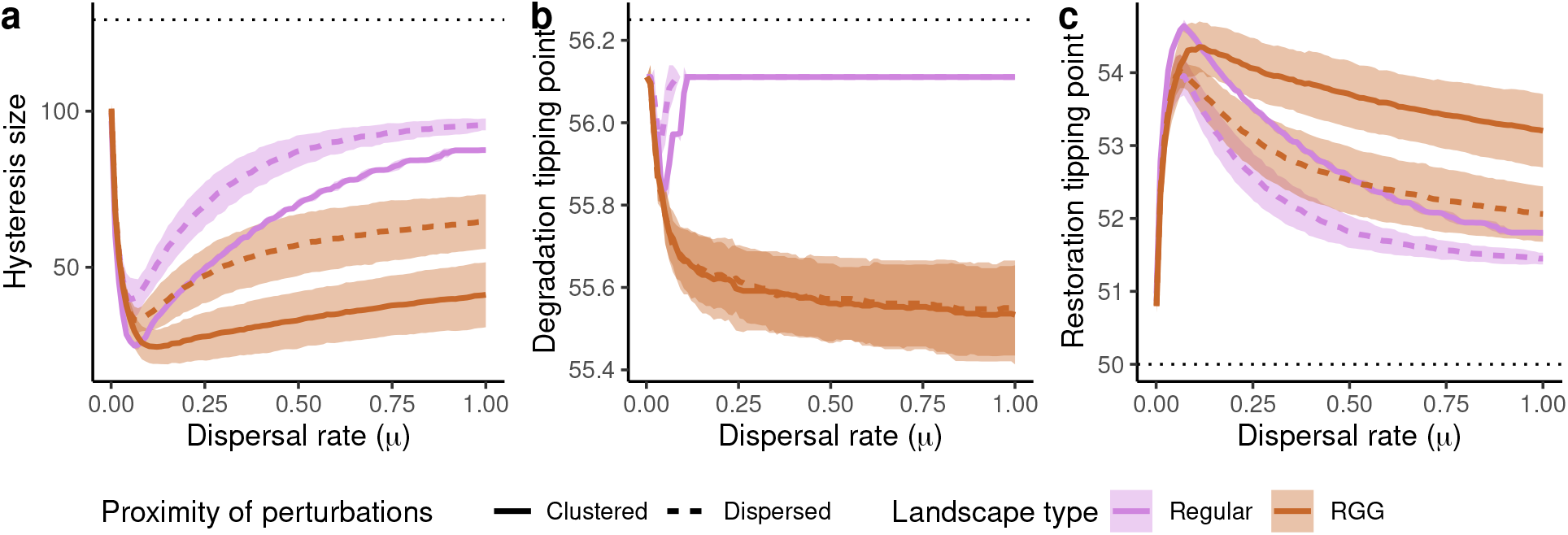
Sensitivity analysis: networks with an average degree of 8. Characterization of the large scale dynamics for different types of perturbations (full line: clustered perturbations, dashed lines: dispersed perturbations) and different spatial structures of metapopulation (purple: regular grid). (a) Size of the hysteresis as a function of the dispersal rate. (b) and (c) large scale tipping points as a function of the dispersal rate: value of the harvesting rate (*B*) at which the system loses (b) or gains (c) the most biomass. All quantities are computed as the mean (lines) and standard deviation of 50 replicates.

**Figure S4.5:**
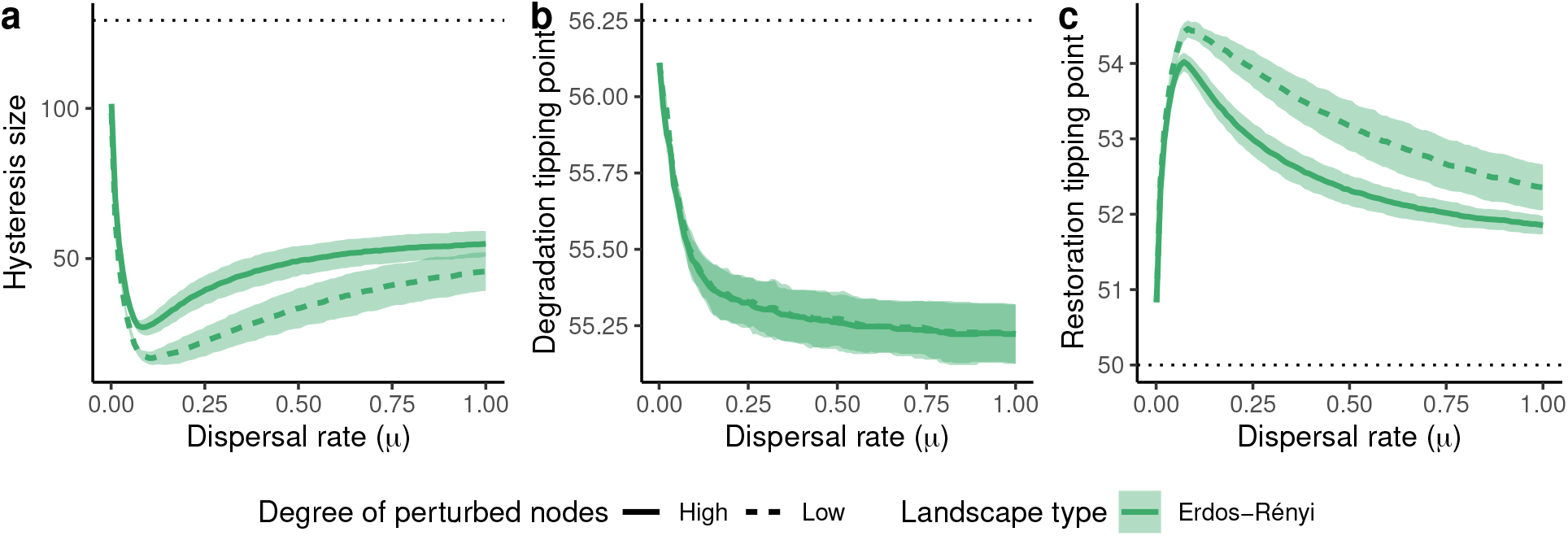
Sensitivity analysis: networks with an average degree of 2. Characterization of the large scale dynamics for different types of perturbations (full line: perturbation of high-degree nodes, dashed lines: perturbation of low-degree nodes) and different spatial structures of metapopulation (green: Erdös-Rényi graphs). (a) Size of the hysteresis as a function of the dispersal rate. (b) and (c) large scale tipping points as a function of the dispersal rate: value of the harvesting rate (*B*) at which the system loses (b) or gains (c) the most biomass. All quantities are computed as the mean (lines) and standard deviation of 50 replicates.

**Figure S4.6:**
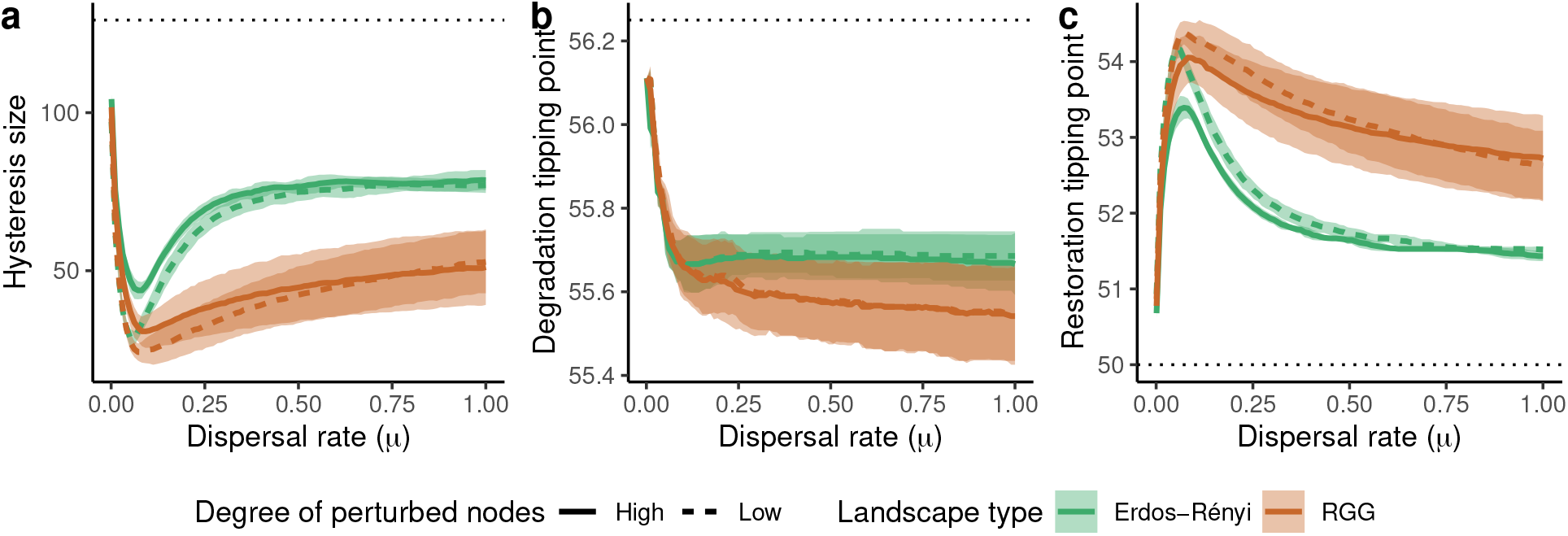
Sensitivity analysis: networks with an average degree of 8. Characterization of the large scale dynamics for different types of perturbations (full line: perturbation of high-degree nodes, dashed lines: perturbation of low-degree nodes) and different spatial structures of metapopulation (green: Erdös-Rényi graphs; brown: random geometric graphs). (a) Size of the hysteresis as a function of the dispersal rate. (b) and (c) large scale tipping points as a function of the dispersal rate: value of the harvesting rate (*B*) at which the system loses (b) or gains (c) the most biomass. All quantities are computed as the mean (lines) and standard deviation of 50 replicates.

### S5 Analysis of a two-patch system

At high dispersal (Fig. S5.1 a, b & c), the densities at equilibrium are the same in the two patches because of the strong homogenization between patches: all equilibria are situated on the *x*_1_ = *x*_2_ plane. Spatial bistability — the stable coexistence of a high- and low-biomass states in the two patches — is not possible, and biomass differences between the two patches can only exist as a transient state towards an homogeneous stable state. The whole system behaves as a single bistable unit with two stable equilibria ((*x*_1_, *x*_2_) = (0,0) and (*x*_1_, *x*_2_) = (*r*_1_, *r*_1_)) separated by an unstable equilibrium ((*x*_1_, *x*_2_) = (*r*_2_, *r*_2_)) over the same range of harvesting rate as for an isolated patch (50 < *B* < 56.25).

As dispersal decreases, the equilibria on the *x*_1_ = *x*_2_ plane keep their values and nature, but branches appear outside of the *x*_1_ = *x*_2_ plane at high harvesting rates (*right before the tipping point toward the desert equilibrium*) (Fig. S5.1 d, e & f). These branches appear from the unstable branch ((*x*_1_,*x*_2_) = (*r*_2_, *r*_2_)) that separates the desert equilibrium ((*x*_1_,*x*_2_) = (0,0)) from the vegetated equilibrium ((*x*_1_,*x*_2_) = (*r*_1_, *r*_1_)) and are themselves unstable. At really low dispersal, spatial bistability becomes possible (meaning the coexistence of patches with different plant biomasses in the metacommunity) since part of the branches outside the *x*_1_ = *x*_2_ plane become stable (Fig. S5.1 g, h & i). These new stable equilibria are associated with strong differences in biomass between the two patches, with one patch at a biomass really close to *r*_1_ while the other is close to 0.

**Figure S5.1:**
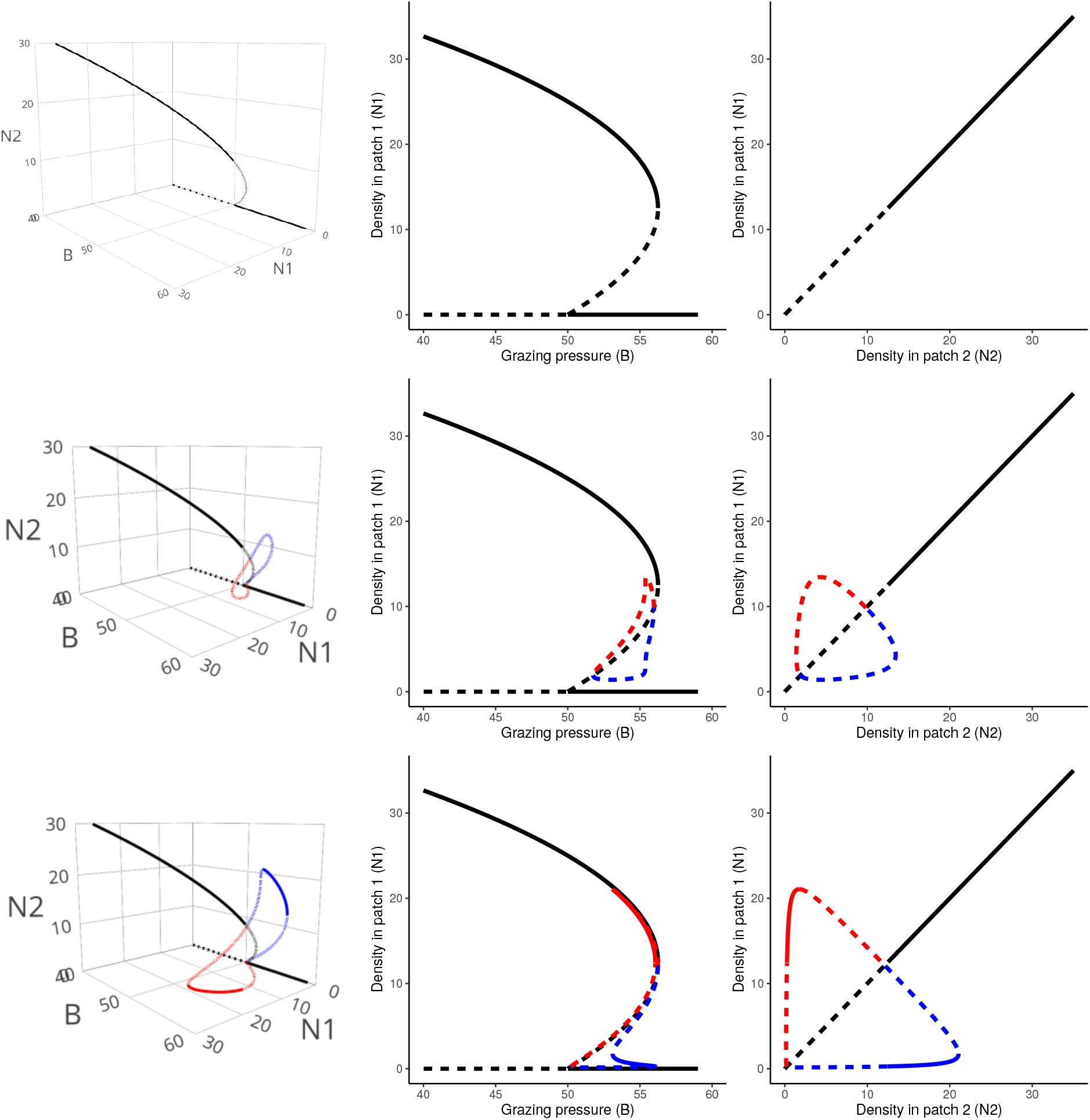
Bifurcation diagrams of a system of two patches connected through dispersal for various dispersal rates: *μ* = 0.3 (top row), *μ* = 0.03 (middle row) and *μ* = 0.005 (bottom row); diagrams in the *n*_1_ − *n*_2_ − *B* space (density in both patches 1 and 2, grazing pressure) in the left column, and their projections on the *B* − *n*_1_ plane (middle column) and *n*_1_ − *n*_2_ plane (right column). Continuous lines depict stable equilibria and dotted lines unstable ones. The branches in black are in the *n*_1_ = *n*_2_ plane (equal densities in both patches), the colored branches are outside of this plane.

### S6 Supplementary figures and tables referenced in the main text

**Table S6.1:** Constant parameters used for the Noy-Meir model.

| Parameter | Value | Biological meaning |
| --- | --- | --- |
| r | 2 | resource growth rate |
| K | 50 | resource carrying capacity |
| A | 25 | half saturation of harvesting |

**Figure S6.1:**
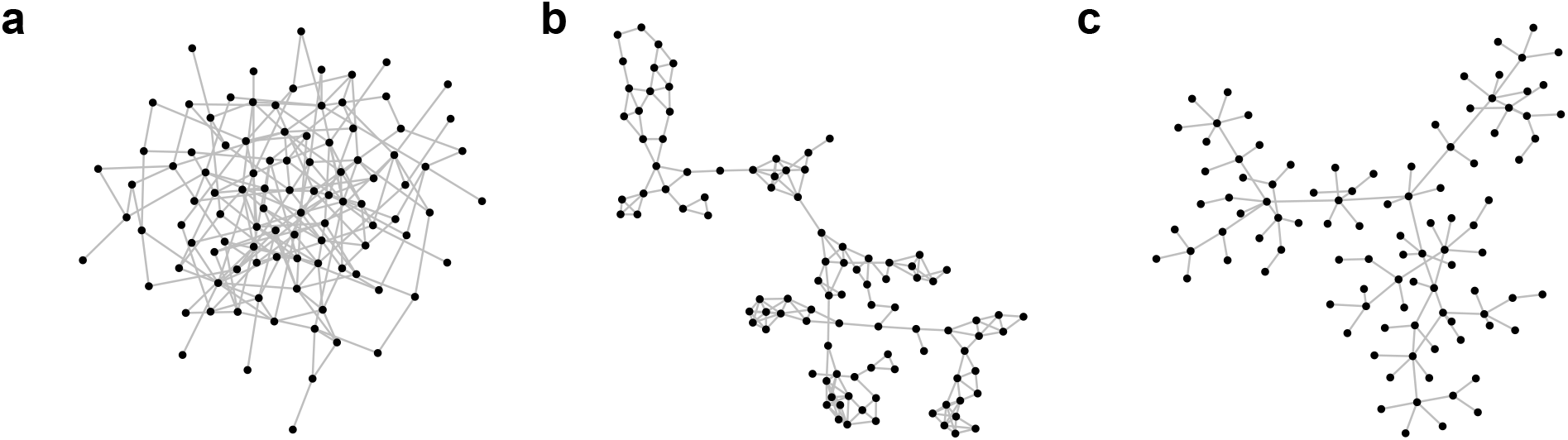
Examples of three landscapes used in the main text analysis. (a) An Erdös-Rényi graph, (b) a random geometric graph (RGG) and (c) an optimal chanel network (OCN).

**Figure S6.2:**
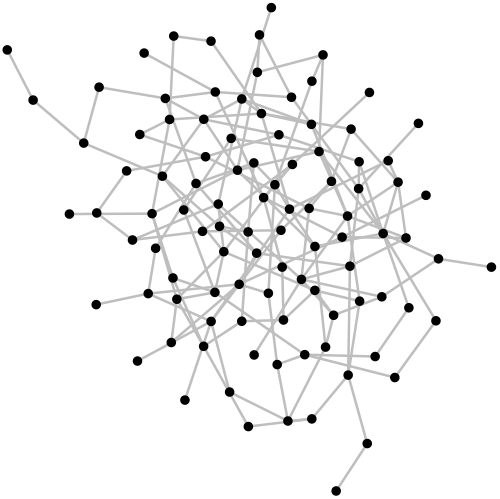
Examples of a low-connectivity Erdös-Rényi graph used in the sensitivity analysis.

**Figure S6.3:**
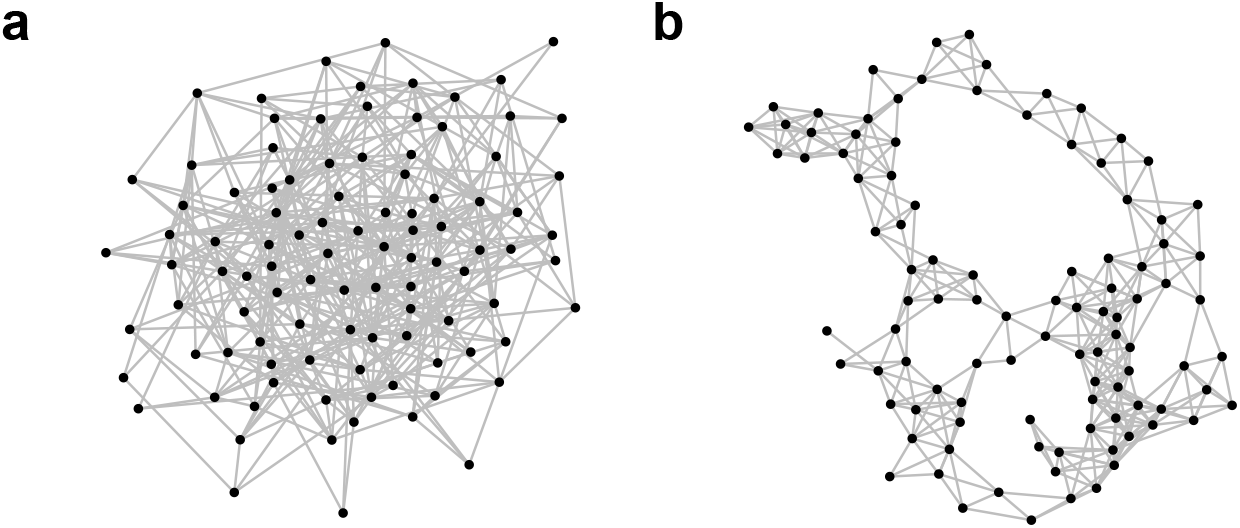
Examples of two high-connectivity landscapes used in the sensitivity analysis. (a) An Erdös-Rényi graph, (b) a random geometric graph (RGG).

